# Tejas functions as a core component of nuage assembly and precursor processing in *Drosophila* piRNA biogenesis

**DOI:** 10.1101/2022.07.12.499660

**Authors:** Yuxuan Lin, Ritsuko Suyama, Shinichi Kawaguchi, Taichiro Iki, Toshie Kai

## Abstract

Piwi-interacting RNAs (piRNAs), a class of 23- to 29-nt gonad-specific small RNAs, function to combat transposons in gonads. piRNAs are thought to be processed and amplified in membrane-less granules called nuage in germline cells. In *Drosophila*, two PIWI family proteins, several Tudor-domain containing (Tdrd) proteins and RNA helicases are assembled at perinuclear region of germline cells, forming nuage to process into piRNAs. Among those, Tejas (Tej), a fly homolog of mouse Tdrd5, has been known as a robust nuage component required for piRNA biogenesis in germline cells, yet its molecular functions remained elusive. To understand its molecular basis on nuage assembly and functions for piRNA biogenesis, we investigated subcellular localization of fluorescent-tagged nuage proteins including Tej and monitored the behavior of piRNA precursors. Tej functions as a core component for assembly of Vasa and Spindle-E to nuage granules through distinct motifs, respectively. The loss of Tej function resulted in malformation of nuage and accumulation of piRNA precursors *en route* in processing, perturbing further piRNA biogenesis in germline cells. Our study also revealed that the low complexity region of Tej regulates the mobility of nuage by phase separation. Collectively, we propose that Tej plays a pivotal role in processing of piRNA precursors by assembling RNA helicases, Vasa and Spindle-E, to nuage, by controlling the dynamics of nuage components.

## INTRODUCTION

Transposons (transposable element, TE) are mobile genetic elements that exist in genomes of all eukaryotic organisms; a substantial part of genomes are indeed occupied by transposons (Samarasinghe et al. 2017; Huang, Burns, and Boeke 2012). They directly impair genomes by causing double-strand breaks, ectopic recombination, and abolishing gene expression (Hedges and Deininger 2007; Hedges and Belancio 2011). PIWI-interacting RNAs (piRNAs), a class of 23- to 29-nt gonad-specific small RNAs function to protect genome integrity, avoiding such catastrophe in germline cells which will be transmitted to next generations (Brennecke et al. 2007; Cenik and Zamore 2011; Czech and Hannon 2011). piRNA is quite conserved and widely found in animals including lowest porifera to primates, and among them, *Drosophila* has been serving as a model system to dissect its molecular mechanisms (Aravin et al. 2006; Girard et al. 2006; Grivna et al. 2006; Hirano et al. 2014; Lim and Kai 2015).

*Drosophila* piRNAs are processed from long piRNA precursor transcripts derived from the genomic loci, called piRNA clusters, where inactive or fragmented transposons are deposited (Brennecke et al. 2007; Brennecke et al. 2008; Malone and Hannon 2009; Khurana et al. 2011; Rozhkov et al. 2013; Ozata et al. 2019). Discrete piRNA clusters on genomic locus are active in gonads, producing dual-strand piRNA precursors in germline cells or uni-strand piRNAs precursors in somatic gonadal cells (Gleason et al. 2018; Ozata et al. 2019). In germline cells, nascent piRNA precursors are transported from the nucleus to the perinuclear region in Nxf3-Nxt1 transport pathway (Kneuss et al. 2019; ElMaghraby et al. 2019; Mendel and Pillai 2019). There, those precursors and transposon RNAs are thought to be processed and/or cleaved in a unique membrane-less structure called nuage, composed of those RNAs, two PIWI family proteins, Aub and Ago3, and other relevant components; DExH-box ATP-binding RNA helicase Vasa (Vas), Spindle-E (Spn-E), and a group of Tudor domain-containing proteins (Tdrds), Krimper (Krimp), Tejas (Tej), Tudor, Tapas, Qin/Kumo, and Vreteno (Gillespie and Berg 1995; Liang, Diehl-Jones, and Lasko 1994; Brennecke et al. 2007; Gunawardane et al. 2007; Golumbeski et al. 1991; Lim and Kai 2007; Patil and Kai 2010; Zamparini et al. 2011; Anand and Kai 2012; Patil et al. 2014). After loading of long piRNA precursors and transposon RNAs to Aub and Ago3, they are sliced into anti-sense and sense piRNAs containing 10-nt complementarity of each other (Aravin et al. 2007; Houwing et al. 2007; Grimson et al. 2008; Lim et al. 2014).

Many of nuage components have Tudor domains that are multifunctional, yet overall activities not fully understood. The Tdrd-family proteins interact with symmetrically-dimethylated arginine (sDMA) residues that are commonly presented in the N-terminus of the PIWI-family proteins (Selenko et al. 2001; Kirino et al. 2009; Li et al. 2009; Kirino et al. 2010). Tudor domain has been shown to promote aggregate formation through sDMA binding in mammalian cells (Courchaine et al. 2021), implying the importance of molecular association by Tdrds for nuage formation. Recent studies have shown that membrane-less organelles composing RNA and proteins are responsible for diverse RNA processing including P-body and Yb body in *Drosophila,* modulating their molecular organization called phase separation (Kistler et al. 2018; Hirakata et al. 2019; Sankaranarayanan et al. 2021). Two of Tdrd proteins localized at *Drosophila* nuage, Tej and Tap, contain an additional conserved Lotus domain that is found in bacteria to eukaryotes (Kubíková et al. 2021). Lotus domain has been demonstrated to interact with the germline-specific DEAD-box RNA helicase, Vas required for piRNA pathway (Jeske, Müller, and Ephrussi 2017).

Of those two, Tej was previously reported as one of the key factors in piRNA pathway both in *Drosophila* and mouse; (Patil and Kai 2010; Yabuta et al. 2011; Patil et al. 2014) A massive reduction of piRNAs and displacement of other components from nuage in the absence of *Drosophila tej* and mouse *Tdrd5* suggest an indispensable role of Tej/Tdrd5 in the piRNA biogenesis, while the molecular functions remained elusive. To explore the detail of the mechanisms of Tej, we re-visited and observed the organization of nuage using fluorescent-tagged nuage components by genome editing and high-resolution microscopy. Loss of interacting domains of Tej with two helicases, Vas and Spn-E, results in failure of proper nuage formation, and concomitantly caused precursor accumulation and TE derepression. We also found that the intrinsically disordered region (IDR) in Tej contributed to the dynamics nuage component. Taken together, our findings propose that Tej plays a pivotal role in engagement of the piRNA precursors to processing by recruiting RNA helicases, Vas and Spn-E, to nuage, and by modulating its dynamics for efficient piRNA biogenesis.

## RESULTS

### Tej associates with Vas and Spn-E, respectively, forming perinuclear nuage granules

We previously reported that Tejas (Tej), a fly homolog of mouse Tudor-domain containing protein 5 (Tdrd5), as a robust piRNA component involved in ping-pong amplification of piRNA in *Drosophila* germline cells (Patil and Kai 2010; Patil et al. 2014). In the absence of *tej*, several components involved in the ping-pong amplification cycle were displaced from nuage, a membrane-less structure where the piRNA amplification takes place, suggesting that general assembly of the nuage requires Tej (Patil and Kai 2010). To dissect the detailed molecular mechanisms involving Tej, we revisited subcellular localization of the nuage components using knock-in fly lines containing fluorescent-tagged proteins generated by genome editing (Gratz et al. 2013). Newly established Tej-GFP, Spn-E-mKate2 (mK2), and mK2-Ago3 as well as previously reported mCherry-Vas, GFP-Vas, and GFP-Aub were examined for their subcellular localization and protein size in ovaries (Fig.1A, S1A, S1B) (Kina et al. 2019). Homozygous fly lines containing *tej^EGFP.KI^, spn-E^mKate2.KI^*, and *ago3^mKate2.KI^*were viable and fertile, suggesting a negligible impact of fluorescent-tag on their functions. Detailed examination of ovaries expressing these nuage component using super-resolution confocal microscope revealed a robust and tight co-localization of Tej with the ping-pong amplification components, two Piwi family proteins, Aub and Ago, and with the RNA helicases, Vas and Spn-E, at perinuclear nuage granules as in those with the antibody staining (Fig.1A, 1B) (Patil and Kai 2010). Consistent with previous immunostainings, localization of Aub and Ago3 were affected in the absence of *tej* (Fig.1B, S1B); fewer GFP-Aub was distributed at the perinuclear region with discernible cytoplasmic fractions, while mK2-Ago3 was observed as cytoplasmic aggregates detached from the perinuclear region (Fig.1B, S1B). Perinuclear nuage granules of both RNA helicases, GFP-Vas and Spn-E-mK2, were also affected in the absence of *tej* (Fig.1B, 1C, S1B). While GFP-Vas was rather found as a smooth layer at the perinuclear region, Spn-E-mK2 was found dominantly in the nucleus of *tej* mutant germline cells. Indeed, co-localization of Vas and Spn-E was perturbed in the absence of *tej* (Fig.1C, S1C).

**Fig. 1.**
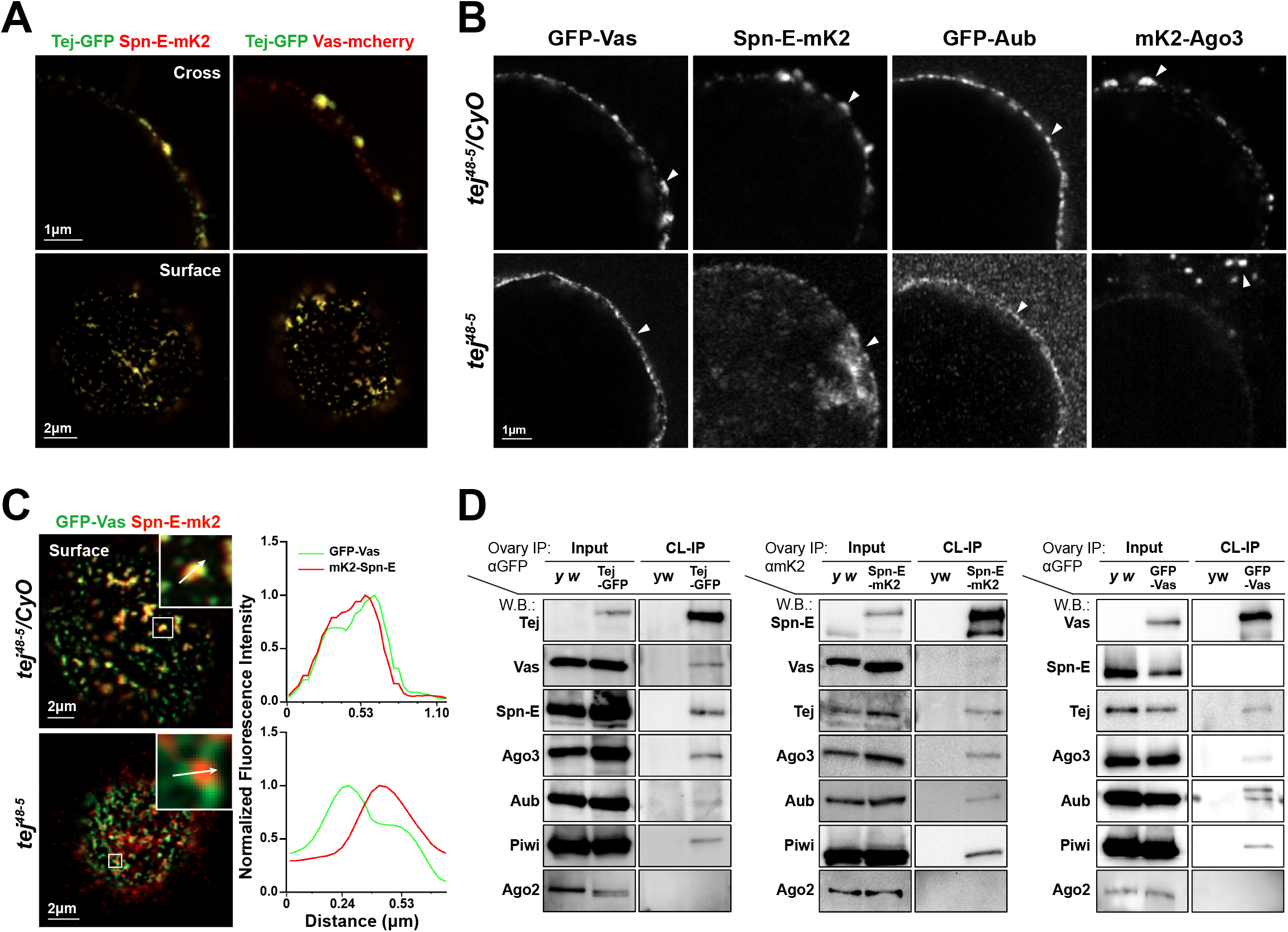
Tej associate with nuage components and is required for proper nuage assembly. **(A)** Co-localization of fluorescent-tagged endogenous Tej-GFP (green) and RNA helicases, Spn-E-mKate2 and Vas-mCherry (red) are shown in the cross-section of the nuclei (top panels) and the surface (bottom panels). **(B, C)** Tej is required for the proper nuage formation. (B) In the control heterozygous, Vas, Spn-E, Aub, and Ago3 are observed as perinuclear granulated puncta (top panels, arrowheads). By contrast, in the absence of *tej*, such perinuclear puncta are perturbed (bottom panels, arrowheads). (C) GFP-Vas (green) and Spn-E-mKate2 (red) are co-localized in the control (top panel) but not in *tej* mutant ovaries (bottom panel). The fluorescence intensity along the designated lines (white arrow in inset) are normalized to the highest value and plotted (right panels). **(D)** Vas and Spn-E mutually exclusively associate with Tej. Immunoprecipitants from ovaries expressing Tej-GFP, GFP-Vas, or mK2-Spn-E are analyzed by western blotting for major piRNA biogenesis factors, Tej, Vas, Spn-E, Ago3, Aub, and Piwi, and an irrelevant siRNA component Ago2.

This segregation of Vas and Spn-E aggregates upon the loss of *tej* prompted us to examine the physical interaction between Tej and Spn-E or Vas *in vivo* by cross-linking immunoprecipitation with Tej (Fig.1D). Using the ovarian lysate expressing tagged nuage components, GFP-Tej was successfully immunoprecipitated with both Spn-E and Vas as well as other major nuage components, Ago3, Aub, and Piwi, but not the irrelevant factor to the piRNA biogenesis, Ago2 (Fig.1D) (Wei et al. 2012). The opposite direction of co-immunoprecipitation using the ovarian lysates expressing either Vas-GFP or Spn-E-mK2 demonstrated their association with Tej and the abovementioned components but not the reciprocal RNA helicase; Vas or Spn-E was hardly detected in immunoprecipitates of Spn-E-mK2 or Vas-GFP, respectively (Fig.1D). These results suggest that Tej interacts with Vas or Spn-E in a different compartment, forming mutually exclusive complexes.

### Tej interact with Vas and Spn-E through the distinct domains

To dissect further the interaction of Tej with Vas and Spn-E, we expressed fluorescent-tagged GFP-Vas, GFP-Spn-E, mK2-Tej in somatic S2 cell line devoid of nuage components or germline-specific factors (Fig.2A, 2B). Upon their single transfection into S2 cells, Vas formed heterogeneous cytoplasmic aggregates in size, while Tej was found as homogeneous ones in the cytoplasm (Fig.2B). By contrast, the majority of Spn-E was dispersed in the nucleus. Interestingly, upon co-expression of full-length Tej (Tej-FL), Vas and Spn-E changed their localization; cytoplasmic Vas and nuclear-localized Spn-E were recruited to large cytoplasmic granules together with Tej, respectively (Fig.2B). These results indicate that Tej can aggregate with cytoplasmic Vas and recruit Spn-E to the cytoplasmic granule.

**Fig. 2.**
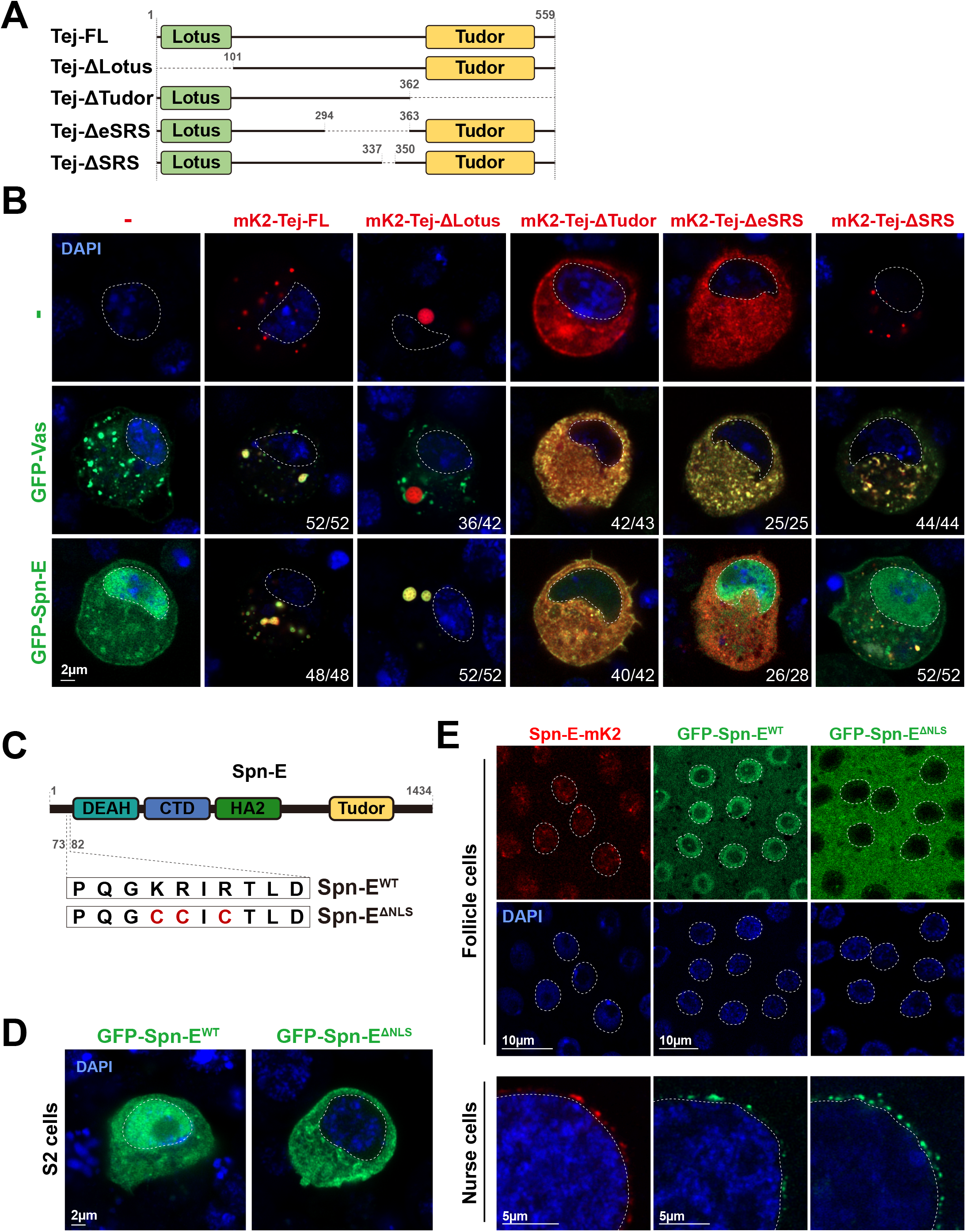
Tejas recruits Vas and Spn-E via individual unique motif. **(A)** Schematic representation of the Tej truncated variants expressed in S2 cells: ΔLotus (deleted 1-100^th^ aa), ΔTudor (deleted 363-559^th^ aa), ΔeSRS (deleted 295-362^nd^ aa), ΔSRS (deleted 338-349^th^ aa). **(B)** Tej recruits Vas and Spn-E via distinct domains. GFP-Vas or GFP-Spn-E (green, middle and bottom panel, respectively) is co-expressed in S2 cells with mKate2-FL-Tej or its truncated variants (red), Tej-ΔLotus, Tej-ΔTudor, Tej-ΔeSRS, and Tej-ΔSRS. The predominant localization of those proteins and the number of cells displaying such patterns are shown. **(C)** Schematic representation of amino acid substitutions in Spn-E NLS (ΔNLS). Lysine at 76^th^ and arginines at 77 and 79^th^ are substituted to cysteines. **(D)** GFP-tagged wild-type Spn-E (Spn-E^WT^, green, left) and NLS-deleted Spn-E (Spn-E^ΔNLS^, green, right) are expressed in S2 cells. **(E)** Spn-E-mK2 shows distinct localizations in ovarian somatic cells and nurse cells (red, left panels). GFP-tagged Spn-E^WT^ or Spn-E^ΔNLS^ is expressed either by a somatic driver, *tj-Gal4,* or a germline driver *NGT40* and *Nos-Gal4*. The nuclei are stained with DAPI (blue) and denoted with dotted circles in (B), (D), and (E).

To identify which domain of Tej is important to recruit Vas and Spn-E, we generated protein truncating variants of Tej and examined their capability for the recruitment (Fig.2A, 2B). Tej harbored two known domains; Lotus domain (6-74^th^ aa) interacts with an RNA helicase, Vas, and Tudor domain (377-488^th^ aa) recognizes a symmetric dimethylation of arginine (sDMA) (Côté and Richard 2005; Jeske, Müller, and Ephrussi 2017). Consistent with the previous observation, Tej devoid of Lotus domain (Tej-ΔLotus) formed cytoplasmic aggregates that failed to co-localize with Vas (Fig.2B). By contrast, Tej-ΔLotus remained colocalized with Spn-E, suggesting that Lotus domain was dispensable for recruiting Spn-E to the cytoplasm and their association (Fig.2B). In contrast, Tudor domain-deleted Tej (Tej-ΔTudor) was dispersed in the cytoplasm with both Vas and Spn-E, suggesting that Tej-ΔTudor is sufficient for the recruitment of Spn-E to S2 cell cytoplasm (Fig.2B). To identify the region required for Spn-E recruitment, we deleted N-terminal part of Tej stepwise and examined for Spn-E localization (Fig.S2A). The control GFP was found both in the nucleus and cytoplasm upon its single transfection, while GFP fused Spn-E was recruited and aggregated in the cytoplasm by co-transfection with Tej variants except for with Tej-Δ1-362 (Fig.S2A). These results indicate that 295-362^nd^ aa is essential to recruit Spn-E to the S2 cell cytoplasm.

Next, we examined whether 295-362^nd^ aa of Tej serves as a motif that recruits Spn-E to the cytoplasm of S2 cells (Fig.2B). Upon co-expression with Tej devoid of 295-362^nd^ aa region, most Spn-E signals remained in the nucleus, while it was dispersed in the cytoplasm (Fig.2B), indicating that Tej 295-362^nd^ aa is a critical part for recruiting Spn-E to the cytoplasm. Alignment of the amino acid sequences among *Drosophila* and the vertebrate homologs showed high conservation for 338-349^th^ aa of *Drosophila* Tej in the proximal region to Tudor domain (Fig.S2B). Deletion of 338-362^nd^ aa (Tej 101-337) from the region of 101-362^nd^ aa between Lotus and Tudor domain significantly impaired Spn-E recruitment to the cytoplasm (Fig.S2C), while deletion of 350-362^nd^ aa (Tej 101-349) did not. In addition, the individual substitutions of amino acids in this region also showed a weaker accumulation of Spn-E in the nucleus, suggesting that 338-349^th^ aa of Tej is critical for the recruitment of Spn-E (Fig.S2C). Thus, we named 338-349^th^ aa region of Tej ‘<u>S</u>pn-E <u>R</u>ecruit <u>S</u>ite’ (SRS) and the 295-362^nd^ as extended SRS (eSRS). Altogether, these results implicate that Tej interacts with Vas and Spn-E via its distinct regions.

Next, we investigated the interacting site of Spn-E with Tej. Based on the predicted ternary structure by AlphaFold v2.1, the C-terminal of the conserved helicase domain of Spn-E was predicted as its potential interaction site to SRS (334-393^rd^ aa; Fig.S2D, S2E) (Jumper et al. 2021). Spn-E containing mutations in the interface region (Spn-E^mut^ ^IN^) was found predominantly in the nucleus when co-expressed with Tej FL (Fig.S2F). As expected, neither Tej-ΔSRS nor -ΔeSRS recruited Spn-E^mut^ ^IN^ (Fig.S2F). Taken together, these results suggest that the residues of Spn-E on the predicted interface with Tej are primarily in charge of the interaction for its recruitment to cytoplasm.

Consistent with the nuclear localization of Spn-E in S2 cells, we found Spn-E had a potential class II monopartite nuclear localization signal (NLS) at the N-terminal part (Fig.2C) (Kosugi et al. 2009). Indeed, point mutations in the core of NLS motif perturbed the nuclear localization of Spn-E in S2 cells (Fig.2D), indicating this sequence served as an intrinsic NLS. Similarly, both fluorescent-tagged endogenous Spn-E-mK2 and transgenic GFP-Spn-E were found in the nucleus of wild-type follicle cells, while they were found in perinuclear nuage in germline cells (Fig.2E). In addition, substituting residues inside the NLS abolished its nuclear localization in the ovarian follicle cells, contrary to its unaffected nuage localization in germline cells (Fig.2E). These results suggest that Spn-E is recruited to the perinuclear nuage in germline cells via germline-specific factor such as Tej, irrespective of NLS signals, exerting an indispensable function for piRNA biogenesis, while its nuclear function remains elusive.

### Tej functions for proper processing of piRNA precursors

Not only the localization of Vas and Spn-E but also that of Aub and Ago3, the PIWI-family proteins required for ping-pong piRNA amplification, have been severely affected in the absence of *tej* (Fig.1B) (Patil and Kai 2010). To further dissect the role of Tej in piRNA biogenesis, 23-29 nt small RNAs bound to Aub or mK2-Ago3 were investigated by next-generation sequencing. As expected, the amount of total piRNAs bound to Aub and mK2-Ago3 was massively reduced in *tej* mutant ovaries compared with those in the heterozygous control (Fig.S3A, S3B). Consistently, piRNAs mapping to the genomic regions, *38C* and *42AB*, where germline specific piRNA precursors were actively transcribed, were remarkably reduced in both Aub- and Ago3-bound populations in *tej* mutant ovaries (Fig.3A). In addition, the 1U preference of antisense Aub-bound piRNA and 10A preference of sense Ago3-bound piRNA populations were notably abolished (Fig.3B). These results suggest that precursor processing into mature piRNAs is perturbed in *tej* mutant ovaries. Consistently, cluster transcripts derived from *38C* and *42AB* were highly accumulated in *tej* mutant (Fig.3C). In contrast, *flamenco*, a piRNA precursor derived from ovarian somatic cell piRNA cluster, was unaffected. Other components required for ping-pong amplification, *vas*, *spn-E*, and *krimp,* also exhibited similar defects, while *nxf3*, a component involved in the transportation of piRNA precursors from the nucleus to the perinuclear region, did not (Fig.3C). These results suggest that Tej functions upstream and/or during the ping-pong amplification after transcription of precursors, and thereby the loss of Tej may cause their accumulation.

**Fig. 3.**
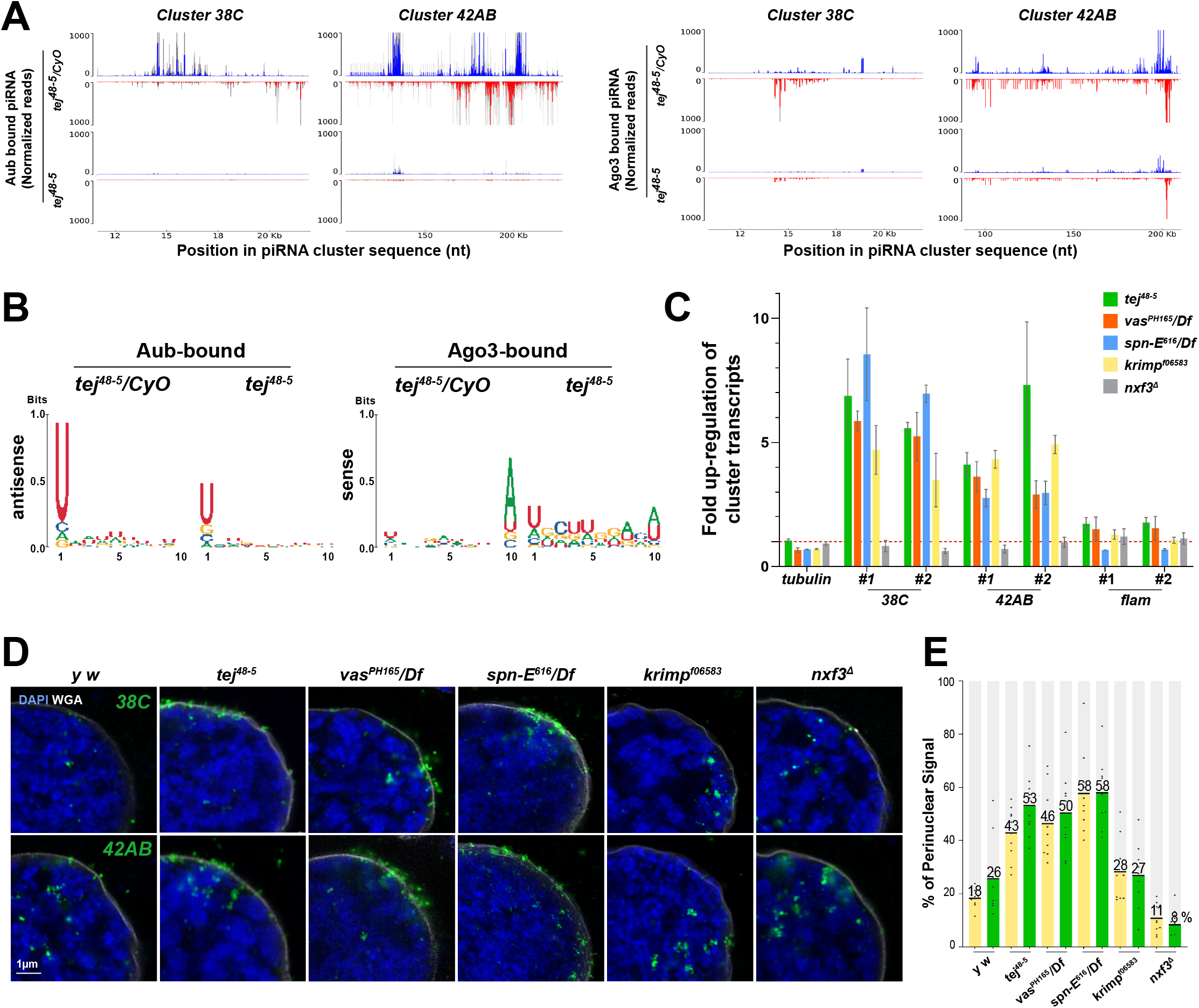
Perturbation of piRNA precursor processing collapses piRNA biogenesis in *tej* mutant ovaries. **(A)** The Aub- (left panels) and Ago3- (right panels) bound small RNAs in the control and *tej* mutant ovaries are mapped to the major piRNA clusters, *38C* and *42AB*. Sense (blue) and antisense (red) piRNAs are indicated with upward and downward peaks, respectively. The grey bars indicate the indpendent biological replicates. **(B)** Nucleotide bias of transposon-mappable Aub- and Ago3-bound piRNAs in the control and *tej* mutant ovaries. piRNA reads are plotted as sequence logos. **(C-E)** Cluster *38C* and *42AB*-derived piRNA precursors are accumulated in the mutant ovaries of piRNA pathway components, *tej*, *spn-E*, *vas*, *krimp,* and *nxf3*. (C) Fold changes of piRNA precursors, c*luster 38C*, *42AB*, and *flam,* in the mutant ovaries. Error bars indicate standard deviation, n=3. (D) piRNA precursors are detected by HCR-FISH (pseudo green) in the control *y w* and the mutant ovaries of indicated genotypes. The nuclear envelope is stained by WGA (pseudo white), and nuclear DNA is stained with DAPI (blue). (E) The ratio of fluorescence intensity of piRNA precursors in the vicinity of nuclear membrane of, *tej*, *spn-E*, *vas*, *krimp,* and *nxf3* mutant germline cells. Signals within 5% both inside and outside of the nuclear membrane are quantified, normalized with the ones in the nucleus (n=10). Numbers on the bars denote percentiles.

To examine the subcellular distribution of piRNA precursors in *tej* mutant, we employed high-resolution RNA-FISH Hairpin Chain Reaction *in situ* Hybridization (HCR-FISH) (Fig.3D) (Choi et al. 2018). Complement probes for cluster transcripts derived from *38C* and *42AB* barely detected their signals as foci mostly in the nucleus of *yw* germline cells (Fig.3D). In contrast, foci were more accumulated in the vicinity of the nuclear membrane, both in the nucleus and cytoplasm of *tej*, *spn-E* and *vas* mutant germline cells (Fig.3D, 3E, mutants; 40-60%, *yw* control;18-26% in average). By contrast, such accumulation was not discernible in *nxf3* mutant, possibly due to failure of their transportation to perinuclear nuage and degradation as previously reported (Fig. 3E, around 10%) (Kneuss et al. 2019). Among those examined piRNA pathway components, *krimp* mutant showed no significant foci in the proximity of perinuclear nuage (Fig.3D, around 28%). These results suggest that Tej may play a role in engagement of piRNA precursors in processing at nuage through the coordination with two RNA helicases, Vas and Spn-E, while Krimp functions during ping-pong amplification, downstream in the processing of precursors (Sato et al. 2015; Webster et al. 2015).

### Tej functions for the processing of piRNA precursors by recruitment of Vas and Spn-E to perinuclear nuage granules *in vivo*

As two distinct motifs, Lotus domain and SRS of Tej, were shown to recruit Vas or Spn-E to Tej granules in S2 cells, respectively (Fig.2B), we next investigated their coordination *in vivo*. Similar to those examined in S2 cells, GFP-tagged Tej variants were expressed with germline drivers, *NGT40*-Gal4 and *nos*-Gal4, in *tej* mutant germline cells, and their localization with mCherry-Vas or Spn-E-mK2 was examined (Fig.4A, 4B). Consistent with our previous study, Tej-FL, as well as Tej-ΔLotus, was observed as granules in the perinuclear region of germline cells (Fig.4B) (Patil and Kai 2010). Tej-FL successfully recruited both Vas and Spn-E onto perinuclear nuage granules, indicating it rescues the proper formation of nuage. In contrast, Vas was significantly segregated from Tej-ΔLotus induced nuage granules and spread as a thin layer in the perinuclear region while Spn-E localization was intact. In addition, although Tej-ΔSRS failed to recruit Spn-E to its aggregates in S2 cells, it did not affect the localization of Spn-E nor Vas at nuage granules *in vivo* (Fig.S4A). These results suggest that deletion of only 12 amino acids of SRS may not be enough to perturb Spn-E recruitment *in vivo*, though piRNA precursors were slightly accumulated, and TE were mildly derepressed (Fig.S4A). To address this possibility, we further deleted an extended region of SRS (Tej-ΔeSRS devoid of 295-362^nd^; Fig.4A, 4B). Tej-ΔeSRS lost its granular formation at the perinuclear region and evenly distributed in the cytoplasm with a lower expression level (Fig.4B, S4B), and failed to restore nuclear accumulation of Spn-E (Fig.4B), indicating that eSRS is essential for adequately recruiting Spn-E to perinuclear nuage granules *in vivo*. However, we cannot exclude a possibility that the lower expression of Tej-ΔeSRS, not deletion of eSRS results in the failure of Spn-E recruitment to nuage granules (Fig.4B, S4B). Tej-ΔTudor lost its granular formation and distributed in both perinuclear and cytoplasmic region. Tej-ΔTudor colocalized partly with Vas and small amount of Spn-E around the perinuclear, although Spn-E was predominantly dispersed in nucleus (Fig.4B), perhaps due to the intact Lotus domain and SRS. In addition, Tej and Spn-E remained in nuage granules in the absence of Vas, but Tej and Vas were displaced from perinuclear nuage without Spn-E (Fig.S4C). These results suggest that Tej and Spn-E function upstream of Vas, while Tej is not simply a hierarchical upstream recruiter of Spn-E due to mislocalization of Spn-E in the absence of Tej, but they are rather mutually dependent for proper assembly into nuage granules. Taken together, our results suggest that each domain of Tej contributes to the proper formation of nuage granules, and both Vas and Spn-E are engaged in this process *in vivo*.

**Fig. 4.**
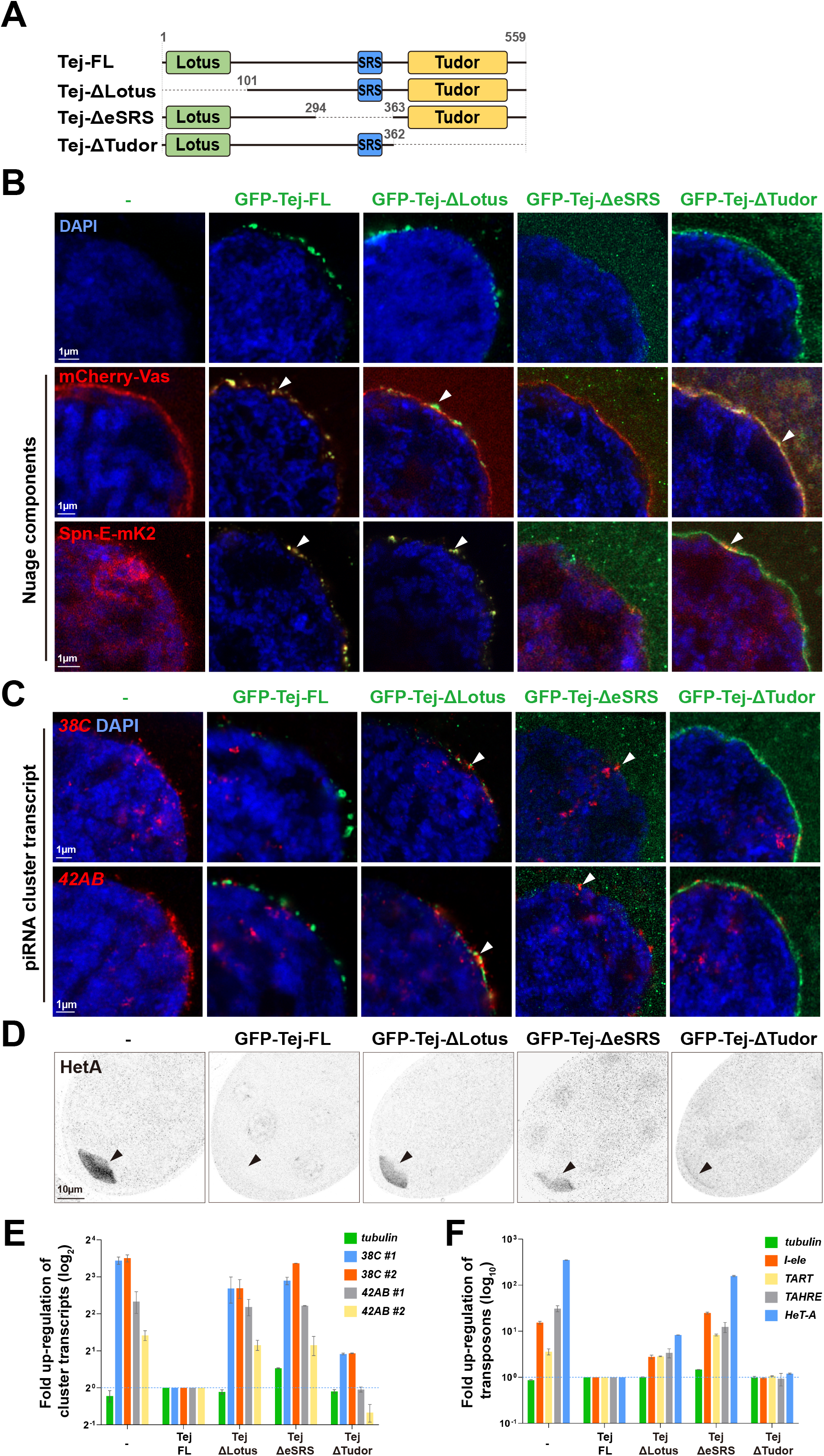
Recruitment of Tej and Spn-E to perinuclear nuage is required for proper processing of piRNA cluster transcripts and TE repression. **(A)** Schematic representation of GFP-Tej truncated variants expressed in germline cells. **(B)** mCherry-Vas (red, middle panels) or Spn-E-mKate2 (red, bottom panels) are observed in *tej* mutant germline cells expressing GFP-tagged transgenes, full-length Tej (Tej-FL), Tej-ΔLotus, Tej-ΔeSRS and Tej-ΔTudor (green, top panels). White arrows indicate the perinuclear aggregations. **(C)** HCR in situ hybridization showing piRNA precursors derived from *cluster 38C* and *42AB* (red) in *tej* mutant germline cells and those expressing Tej variants. White arrows indicate precursors accumulated in the proximity of perinuclear nuage granules. DNA is stained with DAPI (blue) in (B) and (C). **(D)** Immunostaining of Het-A Gap protein in *tej* mutant germline cells and those expressing Tej variants. Black arrows indicate accumulation of Het-A Gag in the oocytes. **(E, F)** The fold changes of transcripts derived from piRNA *cluster 38C* and *42AB* (E) and those of *I-element*, *TART*, *TAHRE*, and *HeT-A* (F) with *tubulin* as a control. All values are normalized to *rp49* and shown as the comparison to the expression level in the presence of Tej-FL. Error bars indicate standard deviation (n=3).

We further examined the accumulation of piRNA cluster transcripts upon the expression of each Tej truncation variant in *tej* mutant germline cells (Fig.4C, 4E). HCR-FISH revealed that the expression of Tej-FL eliminated the accumulation of precursor transcripts derived from the cluster *38C* and *42AB* at the perinuclear region of *tej* mutant germline cells, like those in the control (Fig.4C, 4E, 3D). Notably, germline cells expressing Tej-ΔLotus still represented the accumulation of precursors that were concentrated around imperfectly assembled nuage granules devoid of Vas. In contrast, germline cells expressing Tej-ΔeSRS showed a milder accumulation despite malformation of nuage. By contrast, germline cells expressing Tej-ΔTudor did not exhibit the accumulation of precursors (Fig.4C). The quantification of precursor transcripts by qRT-PCR was also showed that Tej-ΔLotus or Tej-ΔeSRS in *tej* mutant caused robust upregulation of cluster transcripts, but Tej-ΔTudor resulted in a milder upregulation (Fig.4E). These results suggest that Lotus domain and eSRS are important to retain the piRNA precursors in the perinuclear region. Upon the perturbation of Vas or SpnE recruitment to perinuclear nuage granules, the engagement of precursors to processing is seemingly stalled and precursors are accumulated at the perinuclear region.

The transcript level of transposons, such as *Het-A*, *I-element, TAHRE*, and *TART* was also similar to that of piRNA precursors and comparable to the intensity of the Het-A staining in the ovaries (Fig.4D-F). We confirmed Tej-FL in *tej* mutant germline cells almost completely restored the repression of *Het-A* by immunostaining and all transposons examined by qRT-PCR, while Tej-ΔLotus did not (Fig.4D, 4F). Unexpectedly, like Tej-FL, Tej-ΔTudor restored the repression of Het-A as well as other examined transposons, indicating that perinuclear Vas colocalized with Tej-ΔTudor and spotty aggregated Spn-E was adequate for transposon repression. Taken together, our results suggest that Tej functions for the processing of piRNA precursors to repress transposons through proper recruitment of Vas and Spn-E to perinuclear nuage granules.

### The intrinsically disordered region of Tej facilitated LLPS and enhanced the mobility of Vas in nuage

The expression of Tej in S2 cells formed cytoplasmic hollow granules (Fig.S5A, 2B, S2A), prompting us to examine whether Tej is employed in the liquid-liquid phase separation (LLPS). Indeed, the middle part of Tej exhibited a high IUPRED score, further supporting its possibility (Fig.5A) (Dosztányi et al. 2005; Mészáros, Erdős, and Dosztányi 2018). We first investigated the aggregate formation of Tej-Vas-Spn-E in S2 cells. Upon co-transfection of these three components, they formed core-shell granules; Spn-E was concentrated in the center, while Vas and Tej were enriched more at the periphery of these granules (Fig.5B). To further verify whether this unique distribution is representative of the LLPS, we treated the cells with 1,6-hexanediol (1,6-HD), a suppressor of LLPS, by disturbing the weak hydrophobic interactions. Upon the treatment, Vas significantly changed its localization from the periphery to the central part of the granule colocalizing with Spn-E, whereas Tej remained at the periphery (Fig.5B). This susceptibility of the granules to1,6-HD suggests that Tej recruits two helicases, Vas and Spn-E to the aggregates, but being separated from each other by LLPS.

**Fig. 5.**
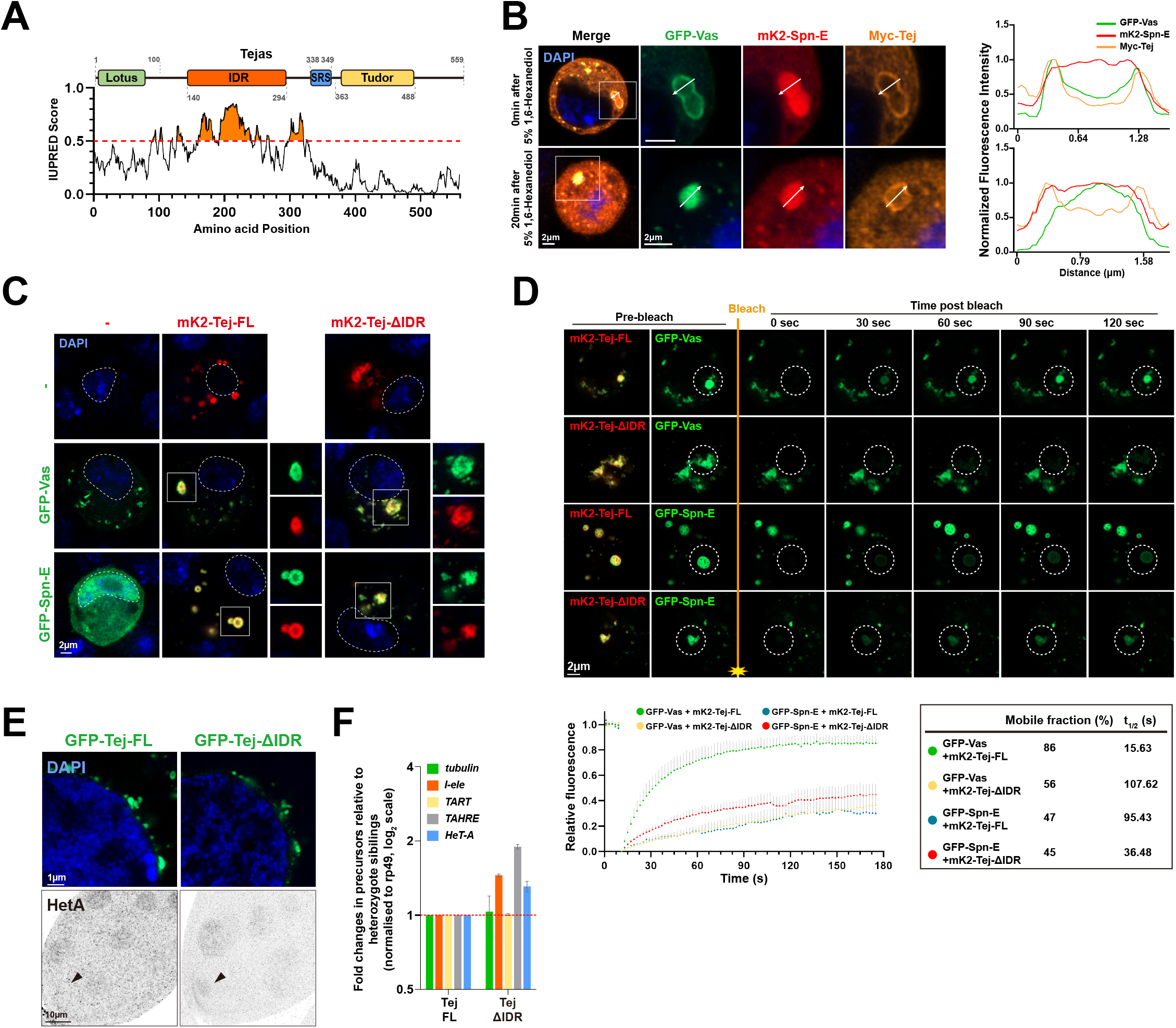
Intrinsically disordered region of Tej controls the mobility of nuage components. **(A)** Schematic structure of Tej intrinsically disordered region (IDR) by the IUPRED prediction. The region with an IUPRED score higher than 0.5 was defined as IDR. **(B)** Tej forms 1,6-hexanediol sensitive core-shell granules with Vas and Spn-E. Myc-Tej is transfected with GFP-Vas and mK2-Spn-E in S2 cells. The fluorescence intensity of each protein is plotted in the diagram with or without 1,6-hexanediol treatment. The intensity profiles of the designated lines (white arrows) are normalized to the highest value of each channel (right panels). **(C)** Recruitment of Vas and Spn-E to Tej condensates is independent of Tej IDR domain. mK2-Tej-FL or Tej ΔIDR is transfected alone (red, top panels) or with GFP-Vas or Spn-E in S2 cells (green, middle and bottom, respectively). Squares indicate aggregates of co-transfected proteins, and each protein is shown in the right panels. **(D)** Tej IDR controls the mobility of Tej aggregates. mK2-Tej FL or Tej ΔIDR (red) is transfected to S2 cells with GFP-Vas or Spn-E (green). Images show the recovery of GFP signals for the aggregates before and after photobleaching (dotted white circles). The line plot shows the normalized relative recovery rate (bottom panel). The mean and ± SD are shown by color dots and gray bars, respectively (n=5). The proportion of mobile fraction and t_1/2_ derived from the mean value of fitting curves are in the table. **(E)** Immunostaining of *tej* mutant ovaries expressing GFP-Tej FL or Tej ΔIDR (top, green) for HeT-A Gap protein (bottom, arrows). **(F) F**old changes of transposon transcripts, *I-element*, *TART*, *TAHRE* and *HeT-A*, and *tubulin* as a control in *tej* mutant ovaries expressing Tej-FL or Tej ΔIDR. All values are normalized to *rp49*, and relative expression levels to those with Tej-FL are shown. Error bars indicate standard deviation (n=3). DNA is stained with DAPI (blue) in (C) and (E).

Next, we further investigated the contribution of domains and the unstructured region of Tej to the formation of cytoplasmic aggregates in S2 cells. Tej-FL formed similar cytoplasmic condensates irrespective of the fluorophores (GFP or mK2) and their location (at N- or C-terminus) (Fig.2B, S2A, S5A), suggesting this aggregation reflects the intrinsic nature of Tej. In addition, Tej-ΔLotus exhibited spherical aggregates, and Tej-Δ1-362 containing Tudor domain alone was sufficient to form amorphous aggregates (Fig.2B, S5A). Meanwhile, the Tej-ΔTudor and Tej-101-362 (101-362^nd^ aa), two Tej variants without Tudor domain, were broadly smeared in the cytoplasm (Fig.2B, S2C, S5A). These results suggest that Tudor domain of Tej is essential for formation of cytoplasmic granules, while the middle IDR part endows their round shape. Notably, Tej-ΔIDR retained the ability to recruit both Vas and Spn-E, respectively (Fig.5C), suggesting IDR region of Tej is dispensable for the recruitment of both RNA helicases.

To examine the contribution of IDR region and its roles in piRNA biogenesis through LLPS, we analyzed the dynamics of Tej, Spn-E, and Vas in S2 cells by fluorescent recovery after photobleaching (FRAP). Upon photobleaching, the fluorescent intensity of GFP-tagged Tej-FL was rapidly recovered in the round granules (Fig.S5B). By contrast, aggregated Tej-ΔIDR and Tej-ΔSRS showed a remarkably and relatively lower recovery rate, respectively. Unexpectedly, Tej-ΔLotus granules were recovered faster than Tej-FL, probably due to the loss of Lotus domain allowing exposure of flexible IDR. These results suggest that Tej IDR facilitates its molecular mobility in S2 cells. Given the high dynamics of Tej granules, we further hypothesized that Tej may contribute to the dynamics of other nuage components in S2 cells. Indeed, upon co-transfection with Tej-FL, more than 80% of Vas was recovered within 90 seconds after photobleaching, while only 56% of Vas was slowly recovered in co-transfection with Tej-ΔIDR, indicating that Tej IDR facilitated the dynamics of Vas (Fig.5D). By contrast, only 45-47% of Spn-E co-expressed with Tej-FL or Tej-ΔIDR was slowly recovered, indicating that Spn-E stayed more stationarily with Tej irrespective of Tej IDR (Fig.5D). Taken together, these results suggest that Tej IDR contributes to Tej mobility and enhances the dynamics of a nuage component Vas, but not that of Spn-E.

Lastly, we investigated a potential function of Tej IDR *in vivo*. *tej* mutant ovaries expressing Tej-ΔIDR formed condensed granules similar to Tej-FL (Fig.5E). However, it showed slight upregulation of Het-A and other transposons by qRT-PCR as well as Het-A expression by immunostaining (Fig.5E, 5F). These results further support the role of IDR for transposon repression, possibly by promoting dynamics of the nuage components for efficient piRNA biogenesis.

## DISCUSSION

In this study, we investigated the molecular role of Tej in nuage assembly with other components and its dynamics that affect the piRNA biogenesis pathway. Visualization of the fluorescent-tagged endogenous nuage proteins, not only PIWI family proteins but also two RNA helicases, Vas and Spn-E, refurbished the defects in their localization in *tej* mutant (Patil and Kai 2010). In addition, piRNA loading to Aub and Ago3 were abolished, and ping-pong amplification was collapsed in *tej* mutant (Fig.3A, 3B). Concomitantly, piRNA precursors were accumulated in the vicinity of nuclear membrane of *tej* mutant germline cells (Fig.3C, 3D). In contrast, *nxf3* mutant that failed to transport piRNA precursors didn’t show any accumulation at the perinuclear region, rather dominantly localization in the nucleus (Kneuss et al. 2019; ElMaghraby et al. 2019). Hence, the accumulation of piRNA precursors at the perinuclear region in *tej* as well as *vas* and *spn-E* mutant germline cells is not relevant to failure of their transport but is likely to be stalled *en route* in processing nearby the malformed nuage (Fig.3D). By contrast, *krimp* mutant did not show such accumulation (Fig.3D), probably because it functions downstream process for further steps of ping-pong amplification cycle, as previously reported for piRNA loading onto Ago3 in OSC cells or recruiting Ago3 to nuage in ovaries (Sato et al. 2015; Webster et al. 2015). Taken together, our results suggest that Tej, Vas, and SpnE may cooperate for the initial phase of the piRNA biogenesis, possibly for the engagement of precursors in processing at nuage (Fig.6).

**Fig. 6.**
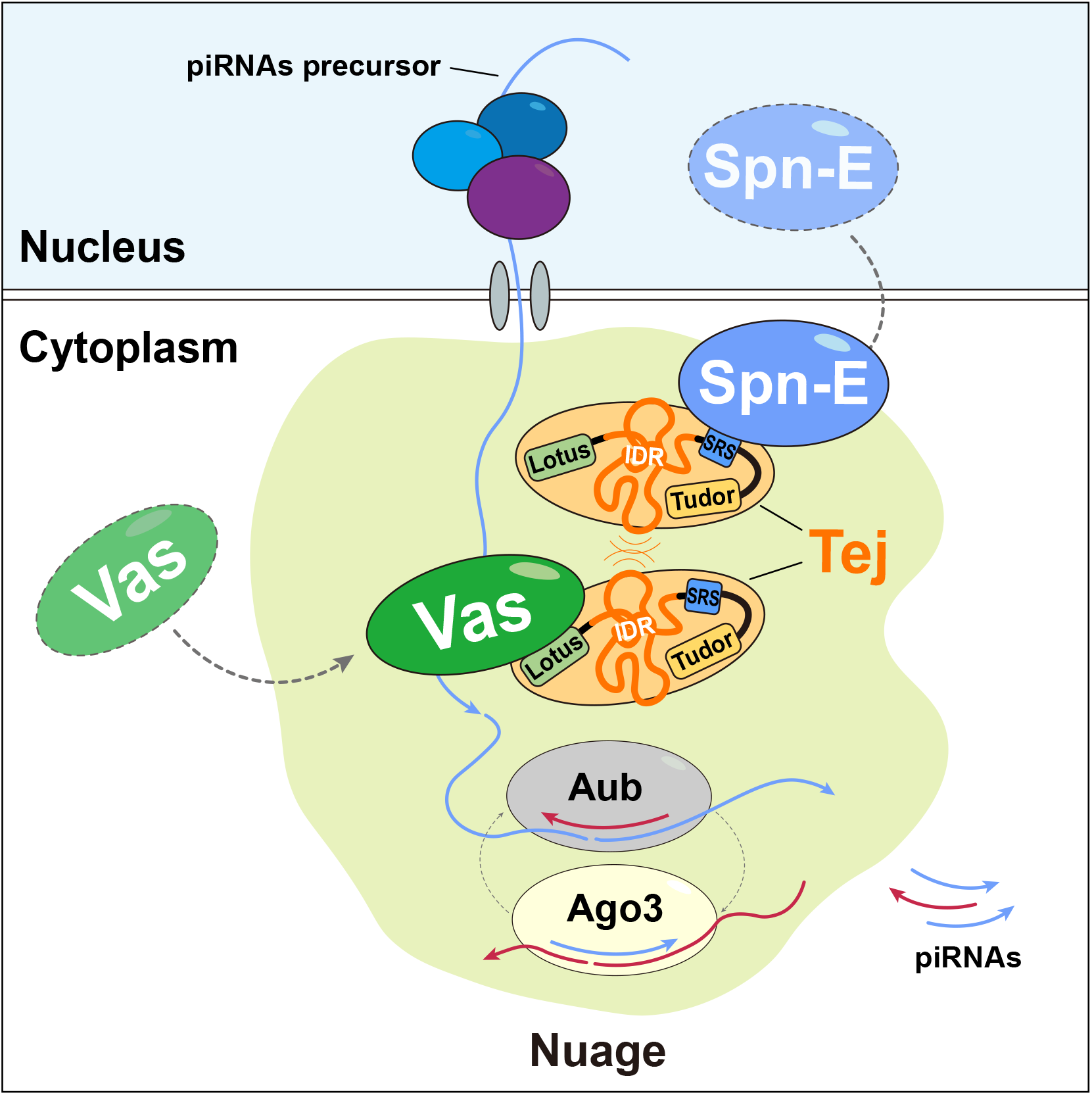
A model: the Tej-mediated nuage organization is essential for the piRNA processing pathway. In the germ cells of *Drosophila*, intrinsically nuclear-localized Spn-E is recruited or intercepted by Tej via its SRS motif and settles together to the cytoplasmic side of the perinuclear region for further nuage organization. Tej is aggregated to nuage granules dependent on the physical interaction or IDR-mediated hydrophobic interaction. Then, through Lotus domain of Tej, Vas is further recruited and employed for the piRNA precursor processing in the nuage. IDR of Tej is allows to modulate the dynamics of other nuage components, especially Vas, in the membrane-less nuage for the efficient piRNA production.

Tej contains two conserved domains, Lotus and Tudor (Patil and Kai 2010; Jeske et al. 2015). Lotus domain is known to stimulate Vas ATP hydrolysis activity required for RNA release (Jeske, Müller, and Ephrussi 2017), implying that Tej may play an important role in adopting Vas to precursor processing during piRNA biogenesis. Indeed, Lotus domain endowed Tej to recruit Vas in S2 cells (Fig.2B, S2A), and piRNA precursors were significantly accumulated around the defective nuage upon deletion of Lotus domain (Fig.4C, 4E). In contrast, Tudor domain that appears in several nuage components was reported to engage in granulation via binding to its ligand, sDMA (Courchaine et al. 2021). Consistently, Tej lacking Tudor domain failed to aggregate by itself but retained the ability of Spn-E recruitment to the cytoplasm in S2 cells (Fig.2B), suggesting that Tudor domain mainly contributes to formation of the condensates (Fig.S5A). However, Tej-ΔTudor suppressed transposon expression to that of the control, despite the aberrant formation of nuage granules and dispersed Spn-E in the nucleus (Fig.4B, 4F). This could be possibly because a few fractions of dispersed Tej-ΔTudor may interact with Vas and Spn-E efficiently nearby nuage through its Lotus and SRS domain, respectively, or make other component associated easily to exert the activity.

Our study revealed that Tej appears to function as a core component for assembly of Vas and Spn-E to nuage granules, via distinct motifs, Lotus and Spn-E Recruit Site (SRS), respectively (Fig.2B, 4B). SRS was responsible for recruitment of Spn-E, which contained a class II monopartite NLS, to cytoplasm out of nucleus (Fig.2C-E). The mouse homolog of Spn-E, TDRD9, was previously reported to be localized in both nuage and nucleus in prespermatogonia (Shoji et al. 2009; Wenda et al. 2017). It raises a possibility that Spn-E as well as its mouse homolog has different functions in the nucleus and nuage, respectively, while their nuclear function remains elusive. Deletion or amino acid substitution of SRS and its predicted interacting site on Spn-E significantly perturbed the recruitment of Spn-E to Tej granules in S2 cells (Fig.2B, S2C-E). However, Tej-ΔSRS almost fully retained Spn-E in perinuclear nuage granules *in vivo* (Fig.S4A), suggesting that components other than Tej and/or other peptide regions in Tej may be involved in tethering Spn-E to the nuage granules *in vivo*. Further deletion of extended SRS (eSRS) resulted in Spn-E localizing in the nucleus *in vivo* as in S2 cells (Fig.4B, 2B), suggesting that Tej without eSRS may have lost such other factors which assist Spn-E recruitment. Nevertheless, we cannot exclude the possibility that the lower expression of Tej-ΔeSRS in germline cells failed perinuclear recruitment of Spn-E (Fig.4B). Our genetic analysis of nuage organization revealed that Spn-E and Tej occupies a higher hierarchical position than Vas and all examined others for the assembly into perinuclear nuage granules (Fig.4B, S4C). It provides a new insight into regulating the stepwise piRNA processing by Tej, Spn-E, and Vas prior to the ping-pong amplification cycle, such as cooperation of Spn-E and Tej leads to precursors processing via their association after capturing the transported long piRNA precursor, and in turn, Vas would function in further processing through Tej Lotus domain or Vas brings transported ones to the SpnE-Tej protein complex (Fig.6). Details of these sequential processes in the complex and the exact feature of precursors processing are still open questions.

As recently reported, LLPS serves to regulate spatial gene expression during *Drosophila* oogenesis; specific RNAs or proteins are modulated in the assembled granules to form the membrane-less structures (Sankaranarayanan and Weil 2020). Osk, a core germplasm protein, interacts with Vas through its Lotus domain, forming the hydrogel structure in the germ plasm (Ephrussi, Dickinson, and Lehmann 1991; Jeske et al. 2015; Kistler et al. 2018). We also found that Tej that has Lotus domain involved in controlling nuage dynamics via Tej IDR. deletion of Tej IDR did not affect the recruitment of Vas and Spn-E, but the shape of their condensates became more amorphous in S2 cells (Fig.5C, S5A). Loss of IDR significantly suppressed the mobility of Tej itself as well as that of Vas in the aggregates (Fig.5D, S5B). In addition, expression of Tej-ΔIDR in *tej* mutant germline cells resulted in a milder upregulation of transposons (Fig.5E, 5F). These results suggest that Tej IDR may function for an efficient piRNA production, potentially by promoting dynamics of nuage components. Interestingly, upon co-transfection of three proteins into S2 cells, Vas and Spn-E were aggregated to Tej granules but segregated into the distinct regions, possibly by LLPS (Fig.5B). DEAD-box RNA helicase family, including Vas, has been previously reported to form non-membranous phase-separated organelles in both prokaryotes and eukaryotes (Hondele et al. 2019), and the large IDR at N terminal moiety facilitates their aggregation by LLPS (Nott et al. 2015). Indeed, our results also showed Vas localization was sensitive to the LLPS suppressor 1,6-HD (Fig.5B), and its mobility was decreased by the loss of Tej IDR (Fig.5D), further supporting LLPS modulate Vas dynamics in the complex. In contrast, Spn-E does not contain notable IDR and forms granules with lesser mobility (Fig.5D). Therefore, Tej may play a central role in a stable complex containing Spn-E and in a mobile one containing Vas in nuage, allowing Vas and Spn-E to compartmentalize with other nuage components via LLPS, as previously reported in the *Bombyx* germ cell line for the stepwise piRNA biogenesis (Nishida et al. 2015). Our co-immunoprecipitation results using ovarian lysate also support this sub-compartmentalization model (Fig1D, Fig.6). IDR of Tej may play a key role not only in engaging precursors in processing together with Vas and Spn-E, but also for ping-pong amplification of piRNAs by controlling proper dynamics of other nuage components, though further investigation would still be required.

## METHODS

### Fly stocks

All stocks were maintained at 25°C with standard methods. The mutant alleles used in the study were *tej^48–5^* (Patil and Kai 2010), *vas^PH165^*(Styhler et al. 1998), *spn-E^616^*(Ott, Nguyen, and Navarro 2014), *krimp^f06583^* (BL# 18990) (Lim and Kai 2007), *ago3^t2/t3^*(BL# 28269; BL# 28270) (Li et al. 2009) *aub^QC42/HN2^*(BL# 4968; BL# 8517) (Schüpbach and Wieschaus 1991), *nxf3^Δ^* (BL# 90328) (Kneuss et al. 2019), *Df(2L)BSC299* (BDSC# 23683), *Df(3R)Exel8162* (BL# 7981). Driver lines for germline and somatic gonadal cells were *NGT40*-Gal4; *nos*-Gal4 VP16 (Grieder, De Cuevas, and Spradling 2000), and *Traffic jam*-Gal4 (DGRC# 104055) (Hayashi et al. 2002), respectively. Either *yw* or the respective heterozygote was used as a control. Knock-In fly lines *vas^mCherry.HA.KI^* (DGRC# 118618), *vas^EGFP.KI^* (DGRC# 118616) and *aub^EGFP.KI^*(DGRC# 118621) (Kina et al. 2019) were obtained from the *Drosophila* Genetic Resource Center at Kyoto Institute of Technology, Japan. All *Drosophila* genotypes used in this study were listed in supplementary table 1.

### Generation of knock-in fly lines

Tej-GFP, mKate2-Ago3, and Spn-E-mKate2 knock-in fly lines were generated through CRISPR-Cas9 induced double-strand breaks restored by the homology-directed repair (HDR) in the presence of donor plasmids. Two guide RNAs were designed to direct the Cas9 proteins to the regions flanking the start/stop codon of each target genes to induce big scale of double-strand breaks. Following guide RNA sequences were cloned into pDCC6 (Gokcezade, Sienski, and Duchek 2014): Tej-GFP gRNA1, GATCGCTCATAGAAACTGGT; Tej-GFP gRNA2, GTGCATAGATTTCTATTATA; mK2-Ago3 gRNA1, TAATAAAAATGCTGGCAATA; mK2-Ago3 gRNA2, TGTGTGTTTCAGAGCATGTC; Spn-E-mK2 gRNA1, GATCACGATGCAATATGGTC; Spn-E-mK2 gRNA2, GAACGATGTAACCATTCTTAT. Donor vectors containing the GFP or mKate2 coding sequence flanked by 1kb homology arms adopted from both 3′ and 5′ sides of the insertion site were generated by cloning of PCR-amplified tags and arms into linearized pGEM-3z vector by In-Fusion HD Cloning Kit (Takara Bio). Obtained gRNA expression plasmids and donor plasmids were injected into the *y w* embryos with a final concentration of 120 ng/ul for each. The Knock-In events positive founders and progenies were confirmed by single fly genome PCR genotyping. Tej-GFP, mKate2-Ago3, and Spn-E-mKate2 Knock-In flies were crossed with corresponding loss-of-function allele *tej^48-5^, Ago3^t2^*, and *Spn-E^616^* for checking the functionality of endogenies fusion proteins, all the fluorescence-fused proteins rescued their corresponding loss-of-function alleles.

### Generation of transgenic fly lines

The transgenic fly lines containing miniTurbo-GFP tagged Tej-FL, Tej-ΔLotus, Tej-ΔTudor, Tej-ΔeSRS, and GFP tagged Tej-ΔSRS, Tej-ΔIDR, Spn-E-FL, Spn-E-ΔNLS were generated by PhiC31 integrase-mediated transgenesis system. The constructs for injection were generated using the cDNAs obtained by reverse transcription from ovarian RNA of *y w* flies. DNA fragments of GFP and the respective variants were amplified and cloned into the pUAS-K10-*attB* plasmid backbone (Koch et al. 2009). The transgenic constructs were injected into the embryo of *attP*-containing strains (P40, BDSC #25709 and P2,BDSC #25710), and progenies expressing mini white were obtained. For rescue experiments, transgenes were recombined with *tej^48-5^* or *Spn-E^616^/Df* background and driven by the germline driver *NGT40-* Gal4; *nos-*Gal4-VP16, or the ovarian somatic cell driver, *traffic jam-*Gal4.

### Antibody generation

Rat anti-Spn-E, Rabbit anti-HeT-A-Gag, rat anti-Ago3 and rat anti-Tej were generated in this study. N terminal GST tagged Spn-E (4-450^th^ aa) antigen peptide was expressed in *E. coli* strain BL21 (DE3) by IPTG with the plasmid generously provided by Dr. M. Siomi. The GST-Spn-E antigen peptide purified by GST affinity beads were used to immunize rats. The Spn-E antibody were further purified from the rat sera with the GST affinity beads conjugated GST-Spn-E antigen peptide and stocked in 50% (v/v) glycerol at −20°C. Plasmid including the fragment that encoding a part of HeT-A-gag (201 amino acids) was generously provided by Dr. Mary-Lou Pardue. DNA fragments encoding N terminal of Tej (1-110^th^ aa) and Ago3 (1-150^th^ aa) were amplified from the cDNA, cloned into pENTR/D-TOPO plasmids, and recombined into either pDEST15 or pDEST17 (Invitrogen). Primers used for the cloning are follows: Het-A gag fw; CACCCCCTACTGGAAAAGCTGAAC, Het-A gag rv; CTACAGGGCATCCTTTGT ACGCGCT, Tej antigen fw; ATGGATGATGGAGGGGAGTT, Tej antigen rv; CTCGGAGGCGTAGCAATA, Ago3 antigen fw: ATGTCTGGAAGAGGAAA, Ago3 antigen rv; TTACACTTCGTAATTAAAAA. The antigens were expressed in *E. coli* strain BL21 (DE3) by IPTG. The purified soluble His-HeT-A-Gag and GST-Tej antigen peptide and the gel pieces of the insoluble GST-Ago3 antigen peptide from SDS-PAGE gel were used to immunize animals (Eve Bioscience, Wakayama). Rabbit serum against HeT-A-gag peptide was directly used for immunostaining. The Tej and Ago3 antibodies were further purified from the sera; insoluble His-Tej antigen and GST-Ago3 antigen peptide-containing region blotted on PVDF membrane (WAKO) was sliced into pieces and incubated with the sera at 4°C overnight with rotation. After incubation, the membrane pieces were washed in 1% (v/v) PBS-Tween for 2h at room temperature, and the antibodies were eluted with 0.1M glycine-HCl (pH 2.5). The elutes were neutralized to pH 7.0 by NaOH and stocked in 50% (v/v) glycerol at −20°C.

### Western blotting

Ovaries were homogenized in the lysis buffer containing 30 mM HEPES (pH 7.4), 80 mM KOAc, 2 mM DTT, 10% (v/v) glycerol, 2 mM MgCl2 and 0.1% (v/v) Triton X-100. After centrifugation at 200,000 ×g for 10 min at 4°C, the supernatants were electrophoresed through pre-cast 5-20% e-PAGEL gels (ATTO) and transferred to ClearTrans SP PVDF membrane (Wako). The primary and secondary antibodies used in this study are listed in supplementary table 2. Antibodies were diluted and stored in the Signal Enhancer reagent HIKARI (NACALAI TESQUE). Chemiluminescence was induced by the Chemi-Lumi One reagent kit (NACALAI TESQUE), and Immunoreactive bands were detected using ChemiDoc Touch (Bio-Rad Laboratories).

### Small RNA immunoprecipitation

For IP of Aub- and mK2-Ago3-bound-piRNAs, 200 ovaries were dissected manually from adult flies in chilled PBS and homogenized with the lysis buffer containing 20 mM Tris-HCl (pH 7.4), 200 mM NaCl, 2 mM DTT, 10% (v/v) glycerol, 2 mM MgCl2, 1% (v/v) Triton X-100, 1x cOmplete protease inhibitor cocktail (Roche) and 1% (v/v) RNaseOUT recombinant ribonuclease inhibitor (Invitrogen). The lysates were cleared by centrifugation at 200,000 ×g for 10 min at 4°C 3 times to remove the contamination of the lipid. Mouse anti-Aub antibody (1:20) (Patil and Kai 2010) or mouse anti-mKate2 (Evrogen, 1:200) was added to the cleared lysate and incubated at 4°C for 2 h with rotation. Then the Dynabeads Protein G/A (Invitrogen) was added to the lysate-antibody mixture and incubated at 4°C for 1 h with rotation. After incubation, the magnet beads were collected and washed at least 4 times with the washing buffer containing 20 mM Tris-HCl (pH7.4), 400 mM NaCl, 2 mM DTT, 10% (v/v) glycerol, 2 mM Mgcl2, 1% (v/v) Triton X-100, 1x cOmplete protease inhibitor cocktail (Roche) and 1% (v/v) RNaseOUT recombinant ribonuclease inhibitor (Invitrogen). 10% of the precipitates were analyzed by western blotting to check the protein immunoprecipitation efficiency. RNAs were isolated from the rest 90% of precipitates with TRIzol LS (Invitrogen) according to the standard manufacturer’s protocol. Purified small RNAs were labeled with ^32^P-γ-ATP using T4 polynucleotide kinase (Thermo Fisher Scientific). After electrophoretic separation by 15% urea-containing denaturing polyacrylamide gel in ×0.5 TBE, radioisotope signals were captured and analyzed by Amersham Typhoon scanner (GE).

### Analysis of small RNA libraries

Small RNA libraries were sequenced using Illumina HiSeq-2500 according to the manufacturer’s protocol at Genome Information Research Center, Research Institute for Microbial Diseases of Osaka University. Small RNA reads were normalized with noncoding RNAs including snoRNAs, snRNAs, miRNAs, and tRNAs. After trimming (5’ adaptor: AGATCGGAAGAGCACACGTCT) and removing rRNA, snoRNAs, snRNAs, miRNAs, and tRNAs, 23- to 29-nt reads were mapped to the piRNA clusters or transposable elements with up to 3-nt mismatching by Bowtie (Langmead et al. 2009). piRNA cluster definition was referred to those previously reported (Brennecke et al. 2007), and TE sequences were adopted from the Flybase (Release 6.32). The normalized numbers of cluster-mapping reads were distributed to the position of the cluster sequence and visualized with pyGenomeTracks (Ramírez et al. 2018). The sequence logos were generated by using ggplot2 R package ggseqlogo (Wagih 2017).

### Crosslinking immunoprecipitation

Ovaries were manually dissected in ice-chilled PBS, fixed with PBS containing 0.1% (w/v) paraformaldehyde for 20 min on ice, quenched in 125 mM glycine for 20 min, and then homogenized in Crosslinking Immunoprecipitation (CLIP) lysis buffer containing 50 mM Tris-HCl (pH 8.5), 150 mM KCl, 5 mM EDTA, 1% (v/v) TritonX-100, 0.1% (w/v) SDS, 0.5 mM DTT and 1x cOmplete protease inhibitor cocktail (Roche). The lysate was incubated at 4°C for 20 min with rotation, followed by 30s sonication with Bioruptor for three times with 30s intervals for cooling (Sonicbio). After centrifugation at 200,000 ×g for 10 min at 4°C, the supernatant was collected in new Eppendorf Protein LoBind tubes and diluted with equal volumes of CLIP wash buffer containing 25 mM Tris-HCl (pH 7.5), 150 mM KCl, 5 mM EDTA, 0.5% (v/v) TritonX-100, 0.5 mM DTT and 1x cOmplete protease inhibitor cocktail (Roche). The diluted lysate was pre-cleaned by Dynabeads Protein G/A (Invitrogen) 1:1 mixture for 1 h at 4°C, and incubated with the antibody (mouse anti-GFP [Thermos Fisher Scientific, 3E6, 1:500] or mouse anti-mKate2 [Evrogen, AB233, 1:500]) overnight at 4°C. Dynabeads Protein G/A (Invitrogen) equilibrated with CLIP washing buffer (1:1 mixture) was added to the lysate-antibody mixture, incubated at 4°C for 3 h. with rotation, collected and washed at least 4 times with the CLIP washing buffer. When required a harsh binding and washing condition, the potassium salt concentration of the CLIP washing buffer was adjusted up to 1M. After washing, beads-bound proteins were retrieved by suspending with the equal volume of the SDS containing 2x sample buffer, heated at 95°C for 5 min, and analyzed through 12% SDS-PAGE gels for Western Blotting.

### RT-qPCR

Total RNAs were extracted from the 2-days old ovaries fattened up with yeast paste with TRIzol LS (Invitrogen) according to the manufacturer’s protocol, treated with DNase I (Invitrogen). cDNAs were generated by reverse transcription with SuperScript III system (Invitrogen) using oligo d(T)20 and hexadeoxyribonucleotide mixture primer. qPCR was performed using KAPA SYBR Fast qPCR Master Mix (KAPA biosystems). All the expression levels of examined genes were normalized to that of *rp49*. The primer sequences for detecting transposon transcripts and piRNA cluster transcripts are shown in supplementary table 3.

### S2 cell culture experiments

*Drosophila* Schneider S2 cells were grown at 26°C in 10% (v/v) Fetal Bovine Serum (FBS) supplemented Schneider medium, with the presence of 50-100 U penicillin and 50-100 µg streptomycin. Plasmids used for transfection were generated using the Gateway cloning system (Life technologies): transgenes were recombined with the *Drosophila* Gateway Vector Collection (DGVC) destination vectors expressing the N terminal tag fused target proteins under Actin5C promoter (Invitrogen, Carlsbad, CA). In addition, a new destination vector for the expression of mKate2 tagged protein at N-terminus under Actin5C promoter, pAKW, was constructed in this study. Transfected S2 cells were placed onto the concanavalin A pre-coated coverslips, incubated at 26°C for at least 20min for an efficient adhesion, fixed for 15 min in 4% (w/v) paraformaldehyde, permeabilized for 10 min in PBX (PBS with 0.2% (v/v) TritonX-100) and washed for 10 min by PBX twice. DNA were stained with DAPI (1:1000) for 10 min and rinsed with PBS. Equilibrated in Fluoro-KEEPER Antifade Reagent (NACALAI TESQUE) for 10min before mounting. Images were taken by ZEISS LSM 900 with Airy Scan 2 using 63× oil NA 1.3 objectives.

### Immunofluorescence staining

Immunostaining of ovaries was conducted as previously reported (Lim et al. 2022). The antibodies used for immunostaining are listed in supplementary table 2. Secondary antibodies were Alexa Fluor 488-, 555-conjugated goat anti-rabbit and anti-mouse IgG (Thermo Fisher Scientific, A11034, A21428, A21127), 1:200 diluted in 0.4% (w/v) BSA containing PBX as working solution. Ovaries expressing endogenous fluorescent-tagged proteins were fixed with PBS containing 0.1% (w/v) paraformaldehyde for 20 min on ice and further washed with PBX (PBS with 0.2% (v/v) TritonX-100) for 10 min twice and washed for 10 min by PBX. DNA were stained with DAPI (1:1000) for 10 min and rinsed with PBS. Equilibrated in Fluoro-KEEPER Antifade Reagent (NACALAI TESQUE) for 10min before mounting. Images were taken by ZEISS LSM 900 with Airy Scan 2 using 63× oil NA 1.3 objectives.

### RNA *in situ* hybridization chain reaction (HCR)

The probes targeting the transcripts derived from the unique regions at *cluster 38C* (Chr2L: 20104896..20213637) and *42AB* (Chr2R: 6322410..6323756) and the reagents were purchased from Molecular Instruments, Inc. The protocol was modified from that was previously reported (Slaidina et al. 2020). Ovaries were fixed in 4% formaldehyde for 20 min, washed twice with PBST at room temperature, and dehydrated by a sequential washing of 25%, 50%, 75%, and 100% (v/v) methanol in PBS for 5 min each on ice. Dehydrated ovaried were stored at −20°C overnight, rehydrated by sequential washes with 100%, 75%, 50%, and 25% (v/v) methanol in PBS on ice. Samples were prewarmed for 2 h in PBX at room temperature, followed by a post-fixation with 4% (w/v) paraformaldehyde, and sequentially washed as follows: twice with PBST for 5 min on ice, once with 50% (v/v) PBST and (v/v) 50% 5× SSCT (5× SSC with 0.1% (v/v) Tween-20) for 5 min on ice, twice with 5× SSCT for 5 min on ice. Then, ovaries were equilibrated with the hybridization buffer for 5 min on ice, prehybridized in the hybridization buffer for 30 min at 37°C and incubated with 0.5 mL of pre-warmed probe hybridization buffer containing 4 pmol of the probes overnight in a light-avoiding 37°C shaker. After hybridization, ovaries were washed 4 times with the probe washing buffer for 15 min each at 37°C and twice with 5× SSCT for 5 min each at room temperature. Next, the ovaries were equilibrated in a prewarmed amplification buffer for 5 min at room temperature. 30 pmol of the probes were denatured at 95°C for 90 sec, chilled down to room temperature for 30 min. Then cool the hairpins on ice for 10 sec and mix with 500 μL amplification buffer at room temperature. The chain reaction was conducted by incubating ovaries in the freshly-prepared the probe solution overnight in a light-avoiding container at room temperature, and terminated by washing twice with 5× SSCT for 5 min. Then the samples were washed in 5× SSCT containing DAPI (1:1000) and Alexa Fluor 488-conjugated Wheat Germ Agglutinin (WGA, 5μg/ml, Thermo Fisher Scientific) and with 5× SSCT for 30 min each at room temperature. Ovaries were equilibrated in Fluoro-KEEPER Antifade Reagent (NACALAI TESQUE) at room temperature before mounting (Choi et al. 2018; Slaidina et al. 2020). Images were taken by ZEISS LSM 900 with Airy Scan 2 using 63× oil NA 1.3 objectives.

### Quantification analysis of in situ-HCR signal for piRNA precursors

Each image was processed with ImageJ (Fiji). Fluorescence intensity of cluster *38C* or *42AB* transcripts by HCR-FISH was measured and quantified after background subtraction. The annular region of ± 5% nuclei diameter inside and outside of the nuclear membrane stained by WGA was defined as the perinuclear region.

### Fluorescence Recovery After Photobleaching (FRAP)

Transfected S2 cells were placed in a concanavalin A-precoated multi-well glass-bottom culture chamber (MATSUNAMI) for over 30 min at 26°C. All images were taken at 26°C in the incubation modules advanced ZEISS LSM 900 with Airy Scan 2 using 63× oil NA 1.3 objectives. One single granule that has GFP signals in each cell was repeatedly bleached using a pulse of 488 lasers 50 times within 3 sec, and images were taken every second to record fluorescence intensity. Initial 10 images were acquired to establish the levels of pre-bleach fluorescence. Fluorescent intensity by bleaching in the specific ROI was analyzed with easyFRAP (Rapsomaniki et al. 2012). A full-scale normalization procedure was used to correct differences in bleaching depth among different experiments, and the recovery curves. Mean recovery curves of normalized data were fit to a double term exponential equation for calculation of the half time of full fluorescence recovery, (t_1/2_[s]), and the percentage of maximum fluorescence recovery (Rapsomaniki et al. 2012).

### Protein disorder prediction and conservation analysis

The intrinsically disordered region was analyzed with the IUPRED server (https://iupred2a.elte.hu/). The region containing residues with IUPRED scores more than 0.5 was classified as a prominent intrinsically disordered region (Dosztányi et al. 2005; Mészáros, Erdős, and Dosztányi 2018).

## Acknowledgment

We thank Dr. Mandy Jeske for useful discussion about unpublished data. Dr. Mikiko C. Siomi generous gifts of plasmid that encode Spn-E antigen. Isshiki Wakana for assisting Spn-E knock-In fly generation, and Alisha Chakrabarti for HeT-A antibodies generation. We acknowledge Bloomington Drosophila Stock Centre and Kyoto Stock Center for the fly stocks. We appreciate the insightful discussion and suggestions from all the members of KT’s laboratory.

## Author Contributions

All authors contributed to the experiment design and data analysis, L YX performed all experiments, L YX, SR, and KS performed the computational analyses, L YX, SR, and KT wrote the paper, KT supervised the project.

## Data And Material Availability

NGS data sets have been deposited to the DNA Data Bank of Japan (DDBJ). BioProject Accession: PRJDB13876. All fly strains and antibodies generated for this study are available upon request.

