## Supplemental Figures for "Tejas functions as a core component of nuage assembly and precursor processing in *Drosophila* piRNA biogenesis"

### Supplementary Figure 1

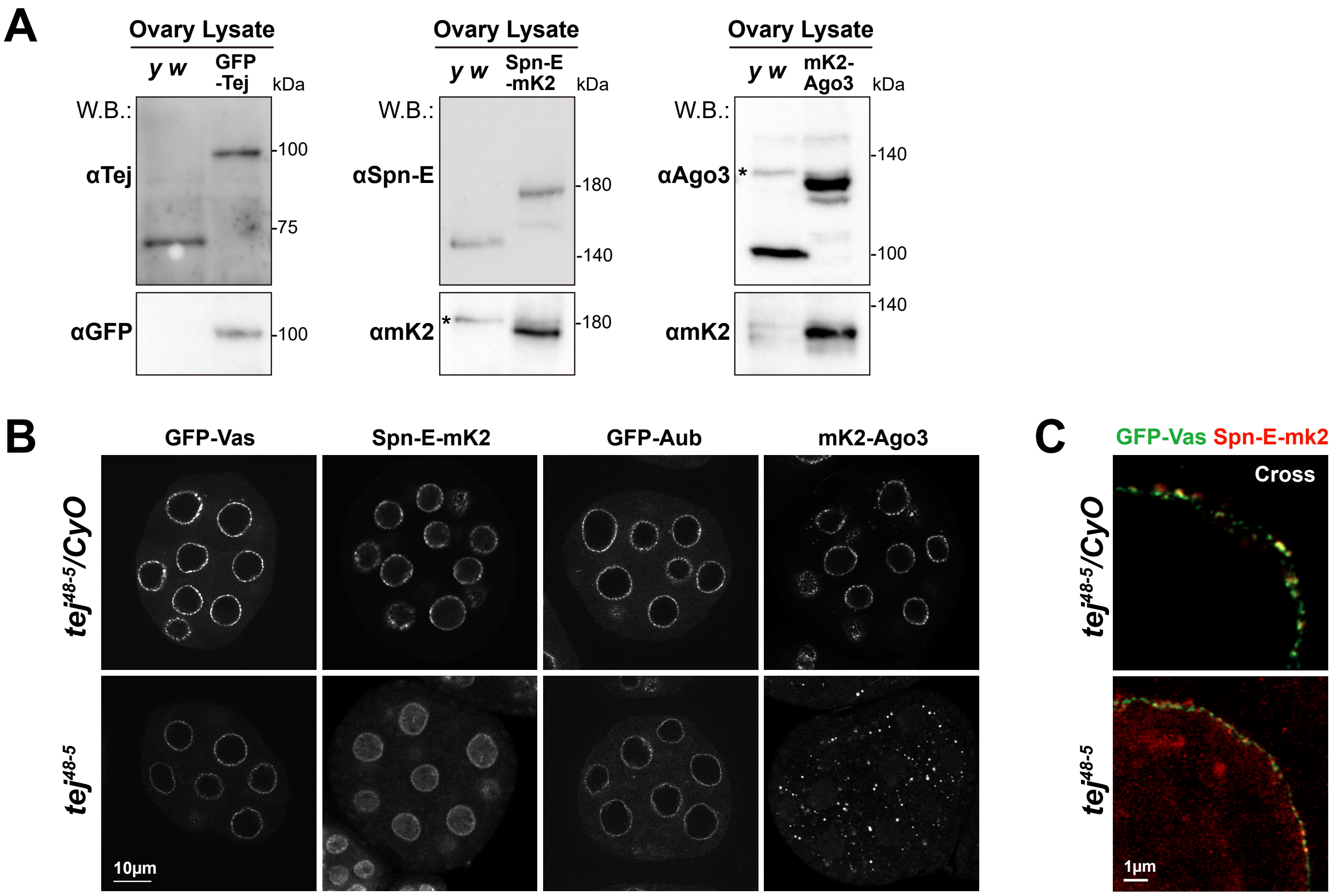

### Supplementary Figure 2

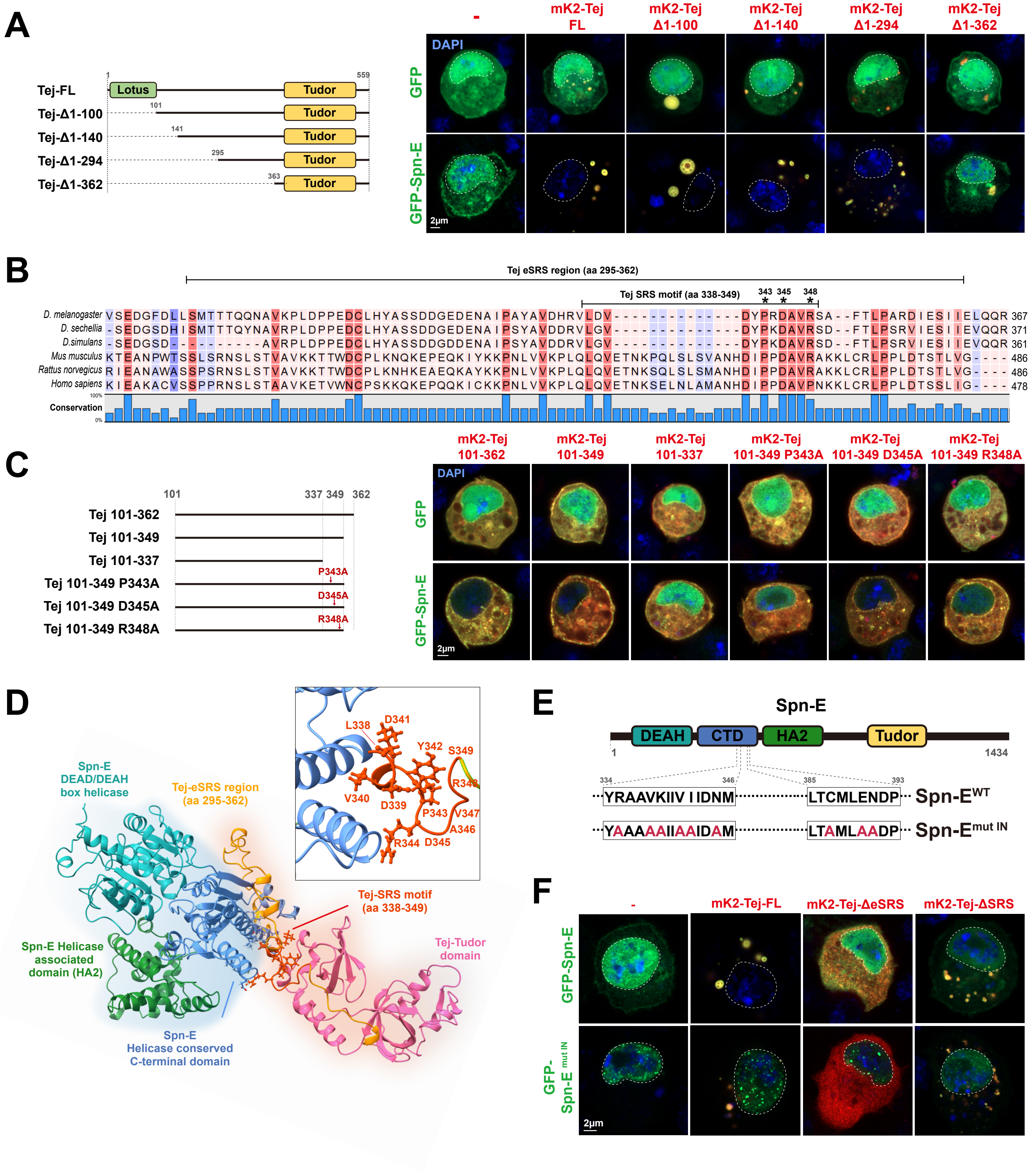

### Supplementary Figure 3

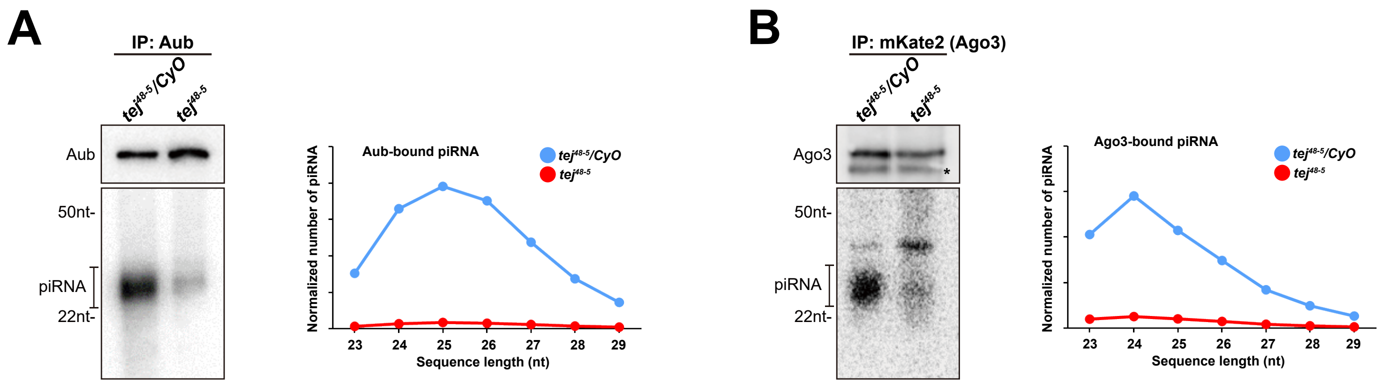

### Supplementary Figure 4

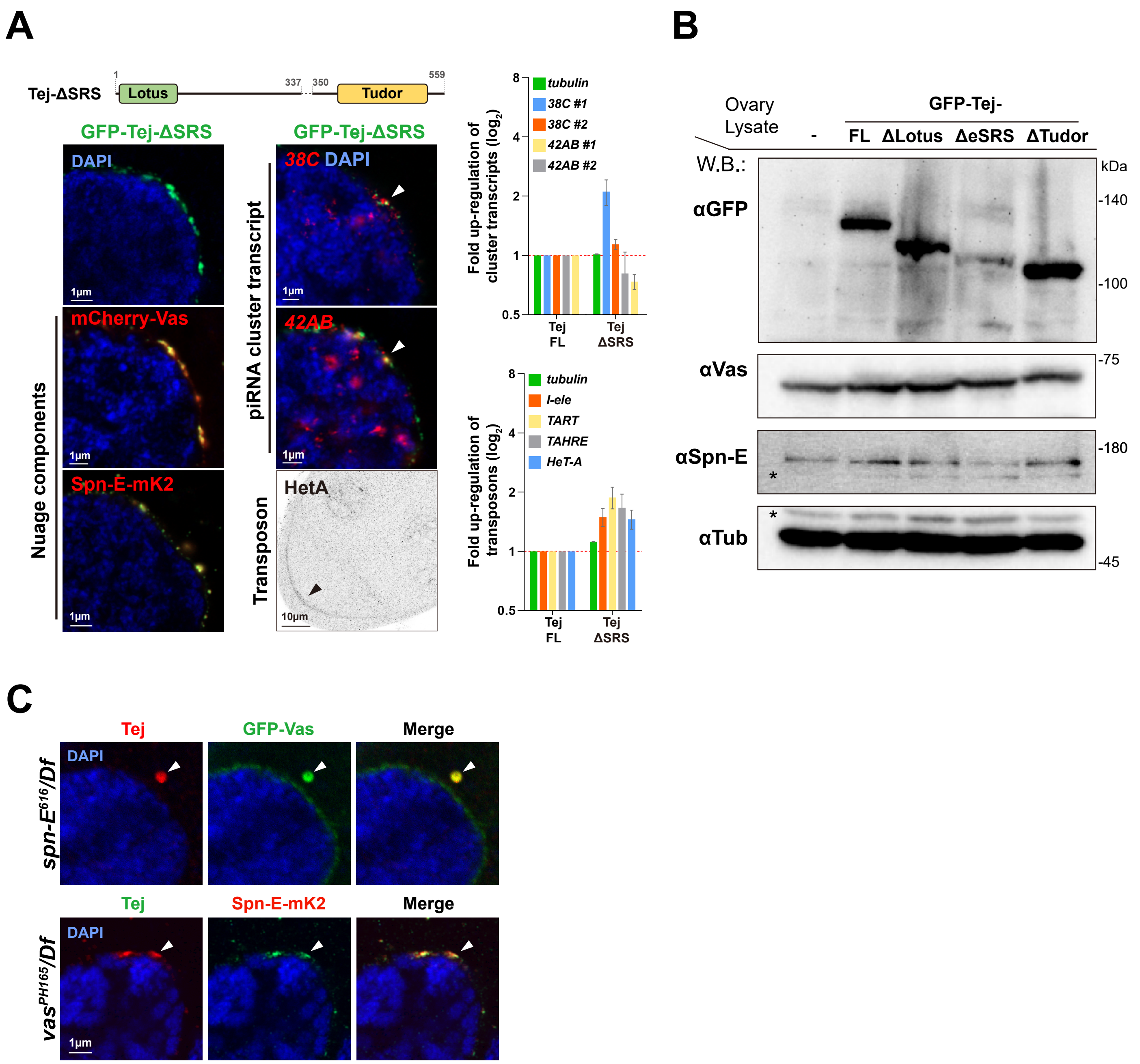

### Supplementary Figure 5

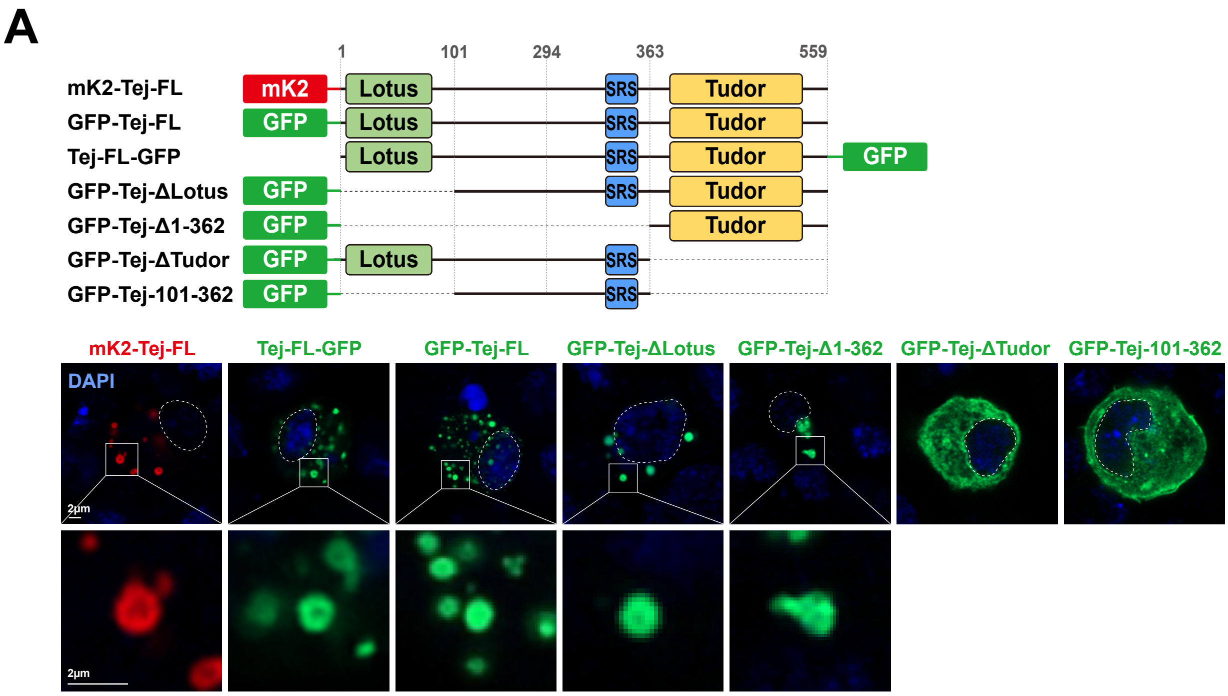


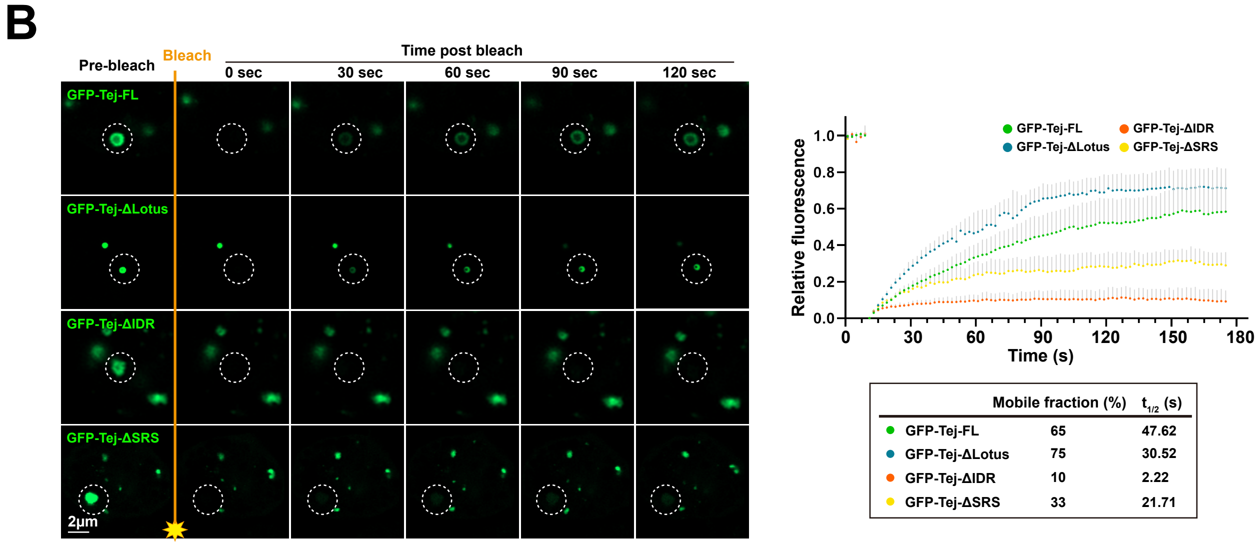


| | Mobile fraction (%) | $t_{1/2}$ (s) |
|---|---|---|
| GFP-Tej-FL | 65 | 47.62 |
| GFP-Tej-ΔLotus | 75 | 30.52 |
| GFP-Tej-ΔIDR | 10 | 2.22 |
| GFP-Tej-ΔSRS | 33 | 21.71 |

**Fig. S1. Endogenously tagged nuage components**

**(A)** Western blot analysis of ovarian lysates showing expression of the newly generated Tej-GFP, Spn-E-mK2, and GFP-Vas by genome editing. Ovaries of *y w* flies are used as a control. Asterisks denote unspecific bands. **(B)** Images of the entire stage 5 egg chambers. Tej is required for the proper nuage formation. Fluorescent-tagged nuage components are present but their localization is affected in *tej* mutant ovaries. All the immunofluorescence signal is represented in grayscale. **(C)** A cross-section of the nurse cell nucleus for each genotype. Co-localization of GFP-Vas and Spn-E-mK2 is lost when Tej is absent.

**Fig. S2. Subcellular localization of Spn-E is controlled by Spn-E NLS and Tej**

**(A)** Identification of Spn-E recruiting domain in Tej. Tej truncated variants are in the schematic representation in left: full-length Tej and Tej devoid of the first 100, 140, 294 and 362 are denoted as FL, Δ1-100, Δ1-140, Δ1-294, and Δ1-362, respectively. GFP or GFP-Spn-E (green, top and bottom panel, respectively) is co-expressed with mKate2-Tej-FL or its truncated variants (red) in S2 cells. Single transfections of those are shown in the left panels. **(B)** Schematic representation of the highly conserved amino acids in Tej. A bar plot of the consavation ratio in percentile is shown on the bottom. SRS and eSRS regions are underlined, while asterisks mark single amino acid substitutions. **(C)** Truncation analysis to search regions associating with Spn-E. Schematic representation of Tej middle part variants (left); Proline, aspartic acid or arginine at 343, 345 and 348$^{th}$ are mutated to alanine, respecitlvely. Either GFP alone or GFP-tagged Spn-E (green) is co-expressed with the mK2-tagged truncated Tej middle part variants (red). **(D)** Predicted interface between Tej and Spn-E by AlphaFoldv2.1 indicates 91-493th amino acid of Spn-E and 295-559$^{th}$ of Tej. The inset shows an enlarged view of the interface. Spn-E recruiting sequence (SRS) is represented in ball-and-stick model. A part of Spn-E in blue represents those predicted to interact with SRS (red) are mutated in the subsequent experiments. **(E)** Schematic representation of Spn-E$^{mut\ IN}$ containing substituted residues (red) that were predicted to be interface with Tej SRS. **(F)** Mutations at Spn-E interface and deletion of Tej SRS abolish recruiting Spn-E to cytoplasm. GFP-tagged Spn-E or Spn-E$^{mut\ IN}$ (green) is co-expressed with the mK2-tagged Tej-ΔSRS or Tej-eSRS (red) in S2 cells. DNA is stained with DAPI (blue) in (A), (C), and (F).

**Fig. S3. Aub- and Ago3 bound piRNAs are remarkably reduced in *tej* mutant ovaries**

**(A, B)** The piRNAs extracted from the immunoprecipitated Aub (A) and mK2-Ago3 (B) in the control and *tej* mutant ovaries are visualized by $^{32}$P-labeling. The immuno-precipitated Aub or mK2-Ago3 are detected by western blot. An asterisk denotes a non-specific band. Line plots show the abundance of Aub and Ago3 bound piRNA in control (blue) and *tej* mutant (red) ovaries by the nucleotide length. Each read number is normalized to those of small RNAs excluding piRNAs.

**Fig. S4. Ovaries expressing Tej variants and genetic hierarchy among Tej, Spn-E and Vas.**

**(A)** Images of ovaries expressing GFP-Tej-ΔSRS with mCherry-Vas and Spn-E-mKate2 (red, right panels). HCR-FISH showing *cluster 38C* and *42AB* transcripts (red, right panels), and immunostaining showing Het-A Gap protein (right, bottom panel, black arrowhead) in *tej* mutant germline cells expressing Tej-ΔSRS. Bar plot shows fold changes of piRNA *cluster 38C* and *42AB* transcripts (top), and the transposon transcripts, *I-element*, *TART*, *TAHRE*, and *HeT-A* (bottom) with a control *tubulin*. All values are normalized to *rp49* and shown as relative expression level compared to ovaries expressing Tej-FL. Error bars indicate standard deviation (n=3). **(B)** Western blotting showing expression levels of GFP-Tej variants from transgenes, and endogenous Vas, Spn-E and Tubulin in ovaries. Asterisks denote non-specific bands. **(C)** Immunostaining of ovaries expressing Spn-E-mk2 (red) in *vas* mutant, and GFP-Vas (green) in *spn-E* mutant, respectively, for Tej. DNA is stained with DAPI (blue). Arrowheads denote Tej granules containing Vas or Spn-E, respectively.

**Fig. S5. Particular domain of Tej controls the morphology and the mobility of Tej-formed aggregations**

**(A)** Schematic drawings represent the fluorophore fused Tej truncations (top). Those GFP- or mK2-tagged Tej variants are expressed in S2 cells. DNA is stained with DAPI (blue). Enlarged images of a single granule are shown at the bottom. **(B)** Fluorescent recovery after photobleaching of each single granule (dotted circles) in GFP-Tej FL or each indicated variants (green) in S2 cells. The line plot shows the normalized relative fluorescence recovery rate. The mean and ± SD are shown by color dots and gray bars, respectively (n=5). The proportion of mobile fraction and $t_{1/2}$ derived from the mean value of fitting curves are in the table.
