## Supplemental Tables for "Tejas functions as a core component of nuage assembly and precursor processing in *Drosophila* piRNA biogenesis"

**Supplementary Table 1. *Drosophila* genotypes used in this study.**

Fig.1A:

*w[-]; tej[EGFP.KI]/ CyO; spn-E[mKate2.KI]/ TM3, Sb*

*w[-]; tej[EGFP.KI], vas[mCherry.HA.KI]/ CyO*

Fig.1B, S1B:

*w[-]; tej*$^{48-5}$*, vas[EGFP.KI]/ CyO*

*w[-]; tej*$^{48-5}$*, aub[EGFP.KI]/ CyO*

*w[-]; tej*$^{48-5}$*/ CyO*; *spn-E[mKate2.KI]/ TM3, Sb*

*w[-]; tej*$^{48-5}$*/ CyO*; *ago3[mKate2.KI]/ TM3, Sb*

Fig.1C, S1C:

*w[-]; tej*$^{48-5}$*, vas[EGFP.KI]/ CyO*

*w[-]; tej*$^{48-5}$*/ CyO* ; *spn-E[mKate2.KI]/ TM3, Sb*

Fig.1D:

*w[-]; vas[EGFP.KI]/ CyO*

*w[-]; tej[EGFP.KI]/ CyO*

*w[-]; spn-E[mKate2.KI]/ TM3, Sb*

Fig.S1A:

*w[-]; vas[EGFP.KI]/ CyO*

*w[-]; tej[EGFP.KI]/ CyO*

*w[-]; ago3[mKate2.KI]/ TM3, Sb*

Fig.2E:

*w[-]; spn-E[mKate2.KI]/ TM3, Sb*

*w[-]; traffic jam*-Gal4/ *CyO; nos*-Gal4 VP16, *Df(3R)Exel8162*/ *TM3, Sb*

*w[-];* UASp-GFP-Spn-E$^{wt}$/ *CyO*; *spn-E*$^{616}$*/ TM3, Sb*

*w[-];* UASp-GFP-Spn-E$^{\Delta NLS}$/ *CyO; spn-E*$^{616}$*/ TM3, Sb*

Fig.3A, 3B, S3A-B:

*w[-]; tej*$^{48-5}$*/ CyO*

*w[-]; tej*$^{48-5}$*/ CyO* ; *ago3[mKate2.KI]/ TM3, Sb*

Fig.3C-E:

*y[-]w[-]*

*w[-]; tej*$^{48-5}$*/ CyO*

*w[-]; vas*$^{PH165}$*/ CyO*

*w[-]; spn-E*$^{616}$*/ TM3, Sb*

*w[-]; krimp*$^{f06583}$*/ CyO*

*w[-]; nxf3*$^{\Delta}$ */TM6C, Sb1*

*w[-]; Df(2L)BSC299/ CyO*

*w[-]; Df(3R)Exel8162/ TM3, Sb*

Fig.4B-F, S4A, S4B, 5E:

*w[-]; NGT40*-Gal4*; nos*-Gal4 VP16

*w[-]; NGT40*-Gal4*; nos*-Gal4 VP16*, spn-E[mKate2.KI]*

*w[-]; NGT40*-Gal4*, vas[EGFP.KI]; nos*-Gal4 VP16

*w[-]; tej*$^{48-5}$*/ CyO*; UASp-miniTurbo-GFP-Tej-FL

*w[-]; tej*$^{48-5}$/ CyO; UASp-miniTurbo-GFP-Tej-ΔLotus

*w[-]; tej*$^{48-5}$/ *CyO*; UASp-miniTurbo-GFP-Tej-ΔTudor

*w[-]; tej$^{48–5}$/ CyO*; UASp-miniTurbo-GFP-Tej-ΔeSRS
*w[-]; tej$^{48–5}$/ CyO*; UASp-GFP-Tej-ΔSRS
*w[-]; tej$^{48–5}$/ CyO*; UASp-GFP-Tej-ΔIDR

Fig.S4C:

*w[-]; vas$^{PH165}$/ CyO; spn-E[mKate2.KI]/ TM3, Sb*
*w[-]; vas[mCherry.HA.KI]/ CyO; spn-E$^{616}$/ TM3, Sb*
*w[-]; Df(2L)BSC299/ CyO*
*w[-]; Df(3R)Exel8162/ TM3, Sb*

**Supplementary Table 2. List of antibodies used in this study.**

| Antibody Name | Source |
|---|---|
| Anti-Tej Rat polyclonal antibody | Current study |
| Anti-Vas Guinea pig polyclonal antibody | (Patil and Kai 2010) |
| Anti-Spn-E Rat polyclonal antibody | (Patil and Kai 2010) |
| Anti-Ago3 Rat polyclonal antibody | Current study |
| Anti-Piwi mouse monoclonal antibody | (Saito et al. 2006) |
| Anti-Ago2 Guinea pig polyclonal antibody | (Iki, Takami, and Kai 2020) |
| Anti-Myc Mouse monoclonal antibody | Wako, Cat.# 017-21871 |
| Anti-eGFP Mouse monoclonal antibody | Invitrogen, Cat.# A-11120 |
| Anti-mKate2 Mouse monoclonal antibody | Evrogen, Cat.# AB231 |
| Anti-HetA Rabbit polyclonal antibody | Current study |
| Anti-Fibrillarin Rabbit polyclonal antibody | Abcam, Cat.# ab5821 |
| Anti-Aub Guinea pig polyclonal antibody | (Lim, L. X. et al., 2022) |
| HRP-conjugated goat anti-guinea pig | Dako, Cat.# P0141 |
| HRP-conjugated goat anti-rat | Dako, Cat.# P0450 |
| HRP-conjugated goat anti-mouse | BioRad, Cat.# 1706516 |
| HRP-conjugated goat anti-rabbit | BioRad, Cat.# 1706515 |

**Supplementary Table 3. List of primers used for qRT-PCR in this study.**

| Primer label | Primer sequence | Source |
|---|---|---|
| Rp49_Fw | ATGACCATCCGCCCAGCATAC | Current study |
| Rp49_Rv | CTGCATGAGCAGGACCTCCAG | Current study |
| Tubulin_Fw | GGTAACCGTCGAAATCAGTGTT | Current study |
| Tubulin_Rv | TGGCTTTTCTGCTATACGTGTC | Current study |
| 38C #1_Fw | GAGACTTGCCGTTCCTTAG | Current study |
| 38C #1_Rv | CCATCTGGAATGCAAACG | Current study |
| 38C #2_Fw | TCCGTGACGGTTTAGCCCA | Current study |
| 38C #2_Rv | AGGTTTCAAACCTTCCAG | Current study |
| 42AB #1_Fw | CGTCCCAGCCTACCTAGTCA | (ElMaghraby et al. 2019) |
| 42AB #1_Rv | ACTTCCCGGTGAAGACTCCT | (ElMaghraby et al. 2019) |
| 42AB #2_Fw | CGCTGTTGAAAGCAAATTGA | (ElMaghraby et al. 2019) |
| 42AB #2_Rv | GAGACCTTCGCTCCAGTGTC | (ElMaghraby et al. 2019) |
| Flam #1_Fw | ACGCTCAGGAAGGGATTTCA | Current study |
| Flam #1_Rv | AAACATGTCGTCTATCCATC | Current study |
| Flam #2_Fw | TCTCGGATAGAACTCTTCCC | Current study |
| Flam #2_Rv | TTGAACCTGTAGGCTAGGTA | Current study |
| HeT-A_Fw | ACAGATGCCAAGGCTTCAGG | (Piñeyro et al. 2011) |
| HeT-A_Rv | GCCAGCGCATTTCATGC | (Piñeyro et al. 2011) |
| TART_Fw | TTCTATCAACAGGCTGTCCACAGGTT | (Savitsky et al. 2006) |
| TART_Rv | CCTTCGTAGTCGGGTAGGATTATTCGT | (Savitsky et al. 2006) |
| TAHRE_Fw | CTGTTGCACAAAGCCAAGAA | (Chen et al. 2016) |
| TAHRE_Rv | GTTGGTAATGTTCGCGTCCT | (Chen et al. 2016) |
| I-element | TGAAATACGGCATACTGCCCCCA | (Klenov et al. 2011) |
| I-element | GCTGATAGGGAGTCGGAGCAGATA | (Klenov et al. 2011) |
